# Silent travelers: the potential of migratory birds and their ticks to introduce emerging viruses into Switzerland

**DOI:** 10.64898/2026.10.29.762706

**Authors:** Corinne Manuela Gerber, Giulia Zezza, Valeria Guidi, Michele Wyler, Claudia Bachofen, Jakub Kubacki

**Affiliations:** Institute of Virology and Immunology (IVI), Mittelhäusern, Switzerland; Department of Infectious Diseases and Pathobiology (DIP), Vetsuisse Faculty, University of Bern, Bern, Switzerland; Institute of Microbiology, University of Applied Science and Arts of Southern Switzerland (SUPSI), Bellinzona, Switzerland

## Abstract

Switzerland hosts over 400 bird species, many of which are migratory. Through diverse movement patterns, ranging from short distances to almost global migrations, these birds can introduce zoonotic viruses, such as West Nile virus (WNV) or avian influenza virus (AIV), as well as unknown and potentially emerging viruses. Furthermore, they facilitate the global dispersal of infected ectoparasites, particularly ticks, risking the introduction of novel tick-borne diseases. To assess this risk, we sampled 42 bird species (39 migratory, 3 resident) over seven months in 2023 at the Bolle di Magadino nature reserve and the Piano di Magadino area in the canton Ticino (Switzerland). This region, because of its transitional climate, serves as an important resting and breeding ground for migratory birds. In total, 496 oropharyngeal-cloacal swabs, 164 sera, and 22 associated ticks were collected.

While RT-qPCR of the swabs detected no active WNV or AIV infections, ELISA testing identified Orthoflavivirus antibodies in 15 of 164 sera, indicating prior exposure. Metagenomic next-generation sequencing revealed a viral landscape spanning 33 families (29 of which were identified in birds). Within this virome, DNA viruses, predominantly *Circoviridae, Genomoviridae,* and *Parvoviridae*, dominated in abundance but exhibited low diversity. In contrast, the highly diverse RNA virome primarily reflected the birds’ dietary habits and the local habitat at the nature reserve. Notably, we report the first detection of Bole tick virus 4 (BoTV4) in Switzerland, identified in several ticks (*Rhipicephalus turanicus*) associated with long-distance migrants. This member of the newly established *Pestiviridae* family is already established in parts of Asia, Africa, and Europe. This study provides the first active surveillance of the wild bird virome in Switzerland. Our findings establish a framework for the early detection of emerging viruses and underscore the role of migratory birds as dispersal vectors for ticks and novel tick-borne viruses.

## Introduction

Wild birds, with over 11,000 different species globally, represent the most diverse tetrapod clade [1–3]. In Switzerland alone, more than 400 species have been recorded [4]. A key ecological feature is their migratory behaviour, with the African-Eurasian flyway acting as a corridor for over two billion birds during their biannual migration, marking it as one of the most critical migratory routes globally [5,6]. Migration distances vary widely among species, from hundreds to several thousand kilometers, with flight durations lasting from one week to a month [7–10]. During migration, birds cross numerous countries and frequently stop at selected stopover sites to rest and feed, where high bird density favours horizontal transmission of microorganisms within and across species [5,11–13].

Birds are flying endothermic vertebrates with a unique adaptive immune system, characterized by a compact genetic repertoire and mechanisms like gene conversion [14–16]. During migration, they must constantly adapt to new environments and pathogens [6,15–18]. Due to the high energetic demands of long-distance flight, a response to novel antigens may be reduced during active migration [19]. This allows for prolonged asymptomatic virus excretion and facilitates host-virus coevolution [13,14]. This phenomenon has been observed in wild waterfowl, the primary reservoir of avian influenza virus (AIV). In Europe, AIV exhibits winter seasonality as aquatic birds congregate in wetlands and viral particles maintain their infectivity for extended periods in cold aquatic environments [13,20–22]. Furthermore, the introduction of West Nile virus (WNV, official species name Orthoflavivirus nilense) Lineage 2 from endemic regions of sub-Saharan Africa into Hungary in the mid-2000s supports the role of birds as a broader viral reservoir. This virus circulates between birds and mosquitoes, which acquire WNV from viraemic birds and transmit it to other birds or incidental hosts such as humans and horses, causing local cases and outbreaks [13,14,23].

Beyond recognized zoonotic pathogens, a high diversity of unidentified viruses has been detected by metagenomic studies, including non-zoonotic or of unclear clinical relevance [24–26]. Viruses from families such as *Astroviridae, Picornaviridae, Adenoviridae, Circoviridae,* and *Parvovirida*e are frequently sequenced from various bird species [27,28]. Additionally, many sequences likely originate from viruses associated with insects, plants, fungi, or bacteria, reflecting dietary intake or environmental exposure rather than active infection [29]. Migratory birds also passively transport hematophagous ectoparasites, particularly ixodid ticks. As a result, African ticks such as *Hyalomma rufipes* are frequently introduced into Europe and may be infected by zoonotic pathogens, most notably the Crimean-Congo haemorrhagic fever virus (CCHFV) [13,30–33].

Despite the increase in research, metagenomic exploration of avian microbial pathogens has predominantly targeted commercial livestock, primarily poultry-related diseases such as *Salmonella* and enteric disorders like runting-stunting syndrome (RSS) or has been restricted to passive monitoring during mass mortality events [15,26,34–48]. While waterfowl are extensively monitored as the main natural reservoir for AIV, this narrow focus creates a gap in global health surveillance regarding other avian taxa [22]. Consequently, research is heavily skewed towards certain species and pathogens, leaving most avian viromes, particularly passerines (songbirds such as sparrows, finches, or swallows), which represent 60% of avian diversity, largely unexplored [2]. While recent studies in China and Australia have begun to fill this gap, no metagenomic analysis has yet investigated migratory birds in Switzerland [2,24]. This represents a vulnerability in One Health preparedness, as ubiquitous birds frequently interface with urban and agricultural environments, increasing the potential for undetected zoonotic spillovers to humans and domestic animals. The limitations of such studies are largely attributable to practical challenges, ranging from complex legal frameworks and small sample sizes to substantial resources required for viral bioinformatics [49–51]. However, establishing a comprehensive viral landscape is essential to provide the baseline necessary for proactive risk assessment.

To address this gap, we performed next-generation sequencing (NGS) to characterize viral diversity in passerines and their associated ectoparasites at the Bolle di Magadino nature reserve (Ticino, Switzerland). This region, as a wetland of international importance and a major stopover site for migratory birds, is an excellent location for sampling and bird monitoring activities [23,52]. Its strategic position in Ticino, in close proximity to WNV-endemic areas in Northern Italy (such as the Po Valley), makes it a vital surveillance point to assess whether migratory birds facilitate the introduction of emerging viruses into Switzerland as they travel along intercontinental flyways [23]. We also investigated whether a bird’s geographic origin and migratory strategy are key determinants of virome composition. In contrast to established diagnostic tools like RT-qPCR and ELISA, which are designed to detect specific pathogens such as AIV and WNV, the NGS-based approach facilitates an exploratory and unbiased detection of the viral landscape that may otherwise remain undetected [53].

## Materials and Methods

### Sampling sites and collection periods

Sampling was conducted in Piano di Magadino (Ticino, Switzerland), a rural area with suitable habitat for multiple bird species located near sites where WNV-positive FTA cards and *Culex pipiens* mosquitoes were detected in 2022 [23]. The Ramsar site of Bolle di Magadino, a unique wetland and marshy habitat, serves as an important resting place for migratory breeders. The sampling season was divided into 3 periods. The spring period (April to mid-May), with mist nets (1 km total length) primarily at the Ramsar site. The reproduction period (mid-May to the end of June), with nets on private property in Cadenazzo. To minimize disturbance during this sensitive nesting period, the sampling was restricted to 10-14-day intervals from 5 a.m. to noon. Lastly, the post-reproductive period (end of June to end of September) involved nets on private property in Cadenazzo and at a farm in Sant’Antonino. Additionally, field rangers provided samples from birds, particularly hooded crows, captured in various locations across Ticino for population containment plans. Capture and handling followed standard ornithological procedures [54,55].

### Sample collection

Bird captures and sampling procedures were conducted under the authorization and supervision of the Cantonal Veterinary Office of Ticino (Permit No. N-35671-TI35/2023/2023) and adhered to Swiss animal welfare legislation. Once captured, birds were identified by species, sexed, aged (if possible), and marked with metal rings before being sampled and released. To facilitate viral preservation, 496 oropharyngeal-cloacal swabs were collected from birds using a swab (DRYSWAB™ ENT MW151, Medical Wire and Equipment, Corsham, Wiltshire, England) and put in a 1 ml UTM™ tube (Universal Transport Medium, Copan, Brescia, Italia) to preserve viruses. Swabs were stored at 4°C for 24-48 hours, then at -80°C. Blood sampling, not exceeding 1% of the birds’ body weight, was performed from the brachial vein of each individual bird using a 25G 5/8 needle (0.5 × 16 mm, Becton Dickinson, Franklin Lakes, NJ, USA). The blood was collected into a non-heparinized calibrated capillary micropipette 100 µl (Cat. No 2-000-100, DRUMMOND, Broomall, PA, USA) and subsequently transferred to a 1.5 ml collection tube using a 15” aspirator tube (Cat. No 2-000-000, DRUMMOND, Broomall, PA, USA). A total of 164 blood samples were kept at 4 °C for 24-48 hours to allow clotting and afterwards centrifuged at 2000 × g, for 10 min at 4 °C to separate serum from clot. Sera were stored at -20 °C while clotted blood at -80 °C, until further analysis. Additionally, a total of 22 ticks were collected, placed in Eppendorf tubes, and kept in a cooling bag until storage at -80°C (Tab 1).

**Table 1:** Bird species, their migratory behaviour, nutrition and numbers of samples collected in this study.

| Migratory behavior | Bird species | Swab | Serum | Tick | Nutrition |
|---|---|---|---|---|---|
| **Resident birds** | *Corvus corone cornix* | 10 | 1 |  | Omnivores |
|  | *Dendrocopos major* | 1 | 1 |  | Insects, seeds |
|  | *Picus viridis* | 2 | 1 |  | Insects |
| **Partial migrants** | *Carduelis chloris* | 2 | 1 |  | Seeds |
|  | *Cettia cetti* | 1 |  |  | Insects, spiders, worms |
|  | *Falco tinnunculus* | 1 |  |  | Insects, small mammals, birds |
|  | *Fringilla coelebs* | 3 | 2 |  | Insects, Seeds |
|  | *Garrulus glandarius* | 1 |  |  | Omnivores |
|  | *Parus major* | 3 | 1 |  | Insects, seeds |
|  | *Passer italiae* | 2 | 1 |  | Insects, seeds |
|  | *Passer montanus* | 3 | 2 |  | Insects, seeds |
|  | *Sylvia melanocephala* | 1 |  |  | Insects, berries |
|  | *Turdus merula* | 7 | 6 | 1 | Fruits, insects, worms |
| **Short-distance migrants** | *Emberiza schoeniclus* | 4 | 2 |  | Insects, seeds |
|  | *Erithacus rubecula* | 53 | 4 | 2 | Insects, berries, spiders |
|  | *Luscinia svecica cyanecula* | 1 |  |  | Insects, spiders |
|  | *Phylloscopus collybita* | 21 |  |  | Insects, spiders |
|  | *Remiz pendulinus* | 1 |  |  | Insects, seeds |
|  | *Serinus serinus* | 4 |  |  | Seeds |
|  | *Sylvia atricapilla* | 154<sup>a</sup> 143<sup>b</sup> | 76 |  | Insects, berries, spiders |
|  | *Turdus iliacus* | 1 |  | 1 | Insects, berries, worms |
|  | *Turdus philomelos* | 2 | 1 |  | Insects, berries, snails, worms |
| **Intermediate migrants** | *Lymnocryptes minimus* | 1 |  |  | Insects, seeds |
| **Long-distance migrants** | *Acrocephalus arundinaceus* | 25<sup>a</sup> 24<sup>b</sup> | 15 | 2 | Insects |
|  | *Acrocephalus schoenobaenus* | 7<sup>a</sup> 6<sup>b</sup> | 1 |  | Insects |
|  | *Acrocephalus scirpaceus* | 34<sup>a</sup> 32<sup>b</sup> |  | 4 | Insects |
|  | *Curruca communis* | 30<sup>a</sup> 27<sup>b</sup> | 7 | 2 | Insects |
|  | *Curruca curruca* | 5<sup>a</sup> 4<sup>b</sup> | 3 |  | Insects |
|  | *Ficedula hypoleuca* | 5<sup>a</sup> 4<sup>b</sup> |  |  | Insects |
|  | *Hirundo rustica* | 14 | 11 |  | Insects, spiders |
|  | *Jynx torquilla* | 6 | 5 |  | Insects |
|  | *Lanius collurio* | 13 | 4 |  | Insects, small mammals, reptiles |
|  | *Locustella naevia* | 2 |  |  | Insects |
|  | *Luscinia megarhynchos* | 4 |  |  | Insects, spiders |
|  | *Luscinia svecica* | 2 | 1 |  | Insects, spiders |
|  | *Muscicapa striata* | 1 |  |  | Insects, spiders |
|  | *Otus scops* | 1 |  |  | Insects |
|  | *Phoenicurus phoenicurus* | 20 | 8 | 1 | Insects, spiders |
|  | *Phylloscopus trochilus* | 23 |  |  | Insects |
|  | *Saxicola rubetra* | 12 | 3 | 9 | Insects, spiders |
|  | *Sylvia borin* | 12 | 6 |  | Insects |
|  | *Ixobrychus minutus* | 1 | 1 |  | Insects, fish, snails, amphibians |
| **Total** |  | **496<sup>a</sup> 476<sup>b</sup>** | **164** | **22** |  |

<sup>a</sup> Number of sampled birds <sup>b</sup> Number of samples available for NGS.

### Molecular and Serological Analysis

Automated viral RNA extraction from oropharyngeal-cloacal swabs was performed on the Maxwell® RSC platform (Promega Corporation, Madison, WI, USA) using 200 µl of UTM™ as starting material. Purification was carried out using the Maxwell® RSC Viral Total Nucleic Acid Purification Kit AS1330 (Promega Corporation, Madison, WI, USA), following the manufacturer’s instruction, with an elution volume of 50 µl of nuclease-free water. Viral RNA from blood clots was extracted manually using the RNeasy Plus Universal kit (Qiagen, Hilden, Germany) and eluted in 60 µl of RNase-free water. The presence of WNV in extracted RNA was analyzed by RT-qPCR using primers and probes, as previously described [23]. Mengo virus strain vMC0 was added to each sample as an internal control of the extraction process and subsequently analyzed by specific RT-qPCR [56]. All RT-qPCR assays were performed on a 7500 Real-Time PCR System (Applied Biosystems™, Waltham, MA, USA). Sera were tested for neutralizing antibodies against WNV using the INGEZIM West Nile COMPAC kit (Gold Standard Diagnostics, Madrid, Spain) following the manufacturer’s instructions. Optical density was measured at 450 nm with Infinite® F50 plate reader and Magellan™ software (Tecan Group Ltd., Männedorf, Switzerland). The inhibition percentage (IP) was calculated in accordance with the protocol. Samples with IP ≥40% were classified as positive, ≤30% as negative and 30-40% as doubtful. To account for the high level of serological cross-reactivity within the *Orthoflavivirus* genus, samples yielding positive or doubtful results were subjected to confirmatory serum neutralization testing (SNT) [57]. The test was performed in vitro with Vero cells (2 × 10^4^ cells/0.1mL), GMEM medium, sodium bicarbonate 7.5% solution, antibiotics (penicillin and streptomycin), antifungal (Amphotericin B) (all Gibco, ThermoFisher Scientific, Waltham, Massachusetts, USA), West Nile virus (isolate France 2018/7025, titer 10^1.9^) and crystal violet staining. To determine the titer of the neutralizing antibodies, a logarithmic dilution of the sera samples was prepared. However, due to the low volume of the starting serum material (<10uL), to be able to perform the assay in duplicates, samples were diluted 1:32, 1:64, 1:128, 1:256 (S2 Table). Consequently, there was insufficient material to perform neutralization assays for other potentially cross-reactive *Orthoflaviviruses*.

### Birds metagenomic sample processing

#### Swab Pooling

All 496 oropharyngeal-cloacal swab samples collected from 42 bird species were screened by RT-qPCR for AIV and WNV. However, 20 samples were excluded from further analysis due to material limitations, like insufficient sample volume after primary screening. Consequently, 476 samples were available for next-generation sequencing (NGS). First, instead of processing each sample individually, samples were pooled predominantly by species (42). Most pools contained a single species. However, four pools contained two species belonging to the same taxonomic family. Overall, 154 pools were prepared, including pools with four birds (n = 96), three (n = 11), two (n = 12), and individual birds (n = 35). Individual samples were analyzed unpooled for birds from which ticks had been removed and from those with a positive WNV antibody ELISA result (INGEZIM West Nile COMPAC kit, Gold Standard Diagnostics, Madrid, Spain). Finally, 200 µl of medium from each sample was transferred to a 1.5-ml Eppendorf tube. The total volume, ranging from 200 to 800 µl, varied depending on the number of birds (one to four). The Mengovirus strain vMC0 (Mengovirus Extraction Control Kit, bioMérieux, Marcy-l’Étoile, France) served as an exogenous internal control at a ratio of 0.5 µl per 100 µl of medium. This control ensured consistent nucleic acid recovery and monitored potential inhibitors [58]. The tube was shaken for 15 seconds (Vortex Genie 2 Digital, Scientific Industries, Bohemia, NY, USA) and centrifuged at 18,500 × g (14,000 rpm) for 10 minutes (Eppendorf Centrifuge 5427 R, Eppendorf, Hamburg, Germany).

#### Filtration

For the filtration process, the supernatant was collected using a 20G needle (0.90 × 40 mm, B. Braun Sterican 100, Melsungen, Germany) and a 1-ml syringe (B. BRAUN Omnifi-F, Melsungen, Germany). The supernatant was then pressed through a 0.45-µm syringe filter (Puradisc, 13 mm, Whatman GE Healthcare, Chicago, Illinois, USA) into a 2-ml Eppendorf tube. This step has been introduced to remove bacteria and host cells bigger than the filter size [59,60].

#### Nucleic acid isolation and SISPA amplification

To achieve a relative enrichment of viral particles, host and environmental nucleic acids not protected by a viral capsid were removed via nuclease treatment, followed by nucleic acid isolation as described previously [60]. To maximize the recovery of the avian virome, total nucleic acids were isolated using the QIAamp Viral RNA Mini Kit (Qiagen, Hombrechtikon, Switzerland) with modifications such as the omission of the carrier RNA to prevent host-derived background interference and addition of β-mercaptoethanol to a final concentration of 1% in the AVL buffer (Gibco, 50 mM, Thermo Fisher, Waltham, Massachusetts, USA) to eliminate any residual nuclease activity from previous nuclease treatment. To elute DNA and RNA viruses, 30 µl of RNase-free water and 30 µl of Tris-EDTA (TE) buffer (1× molecular biology grade, Thermo Fisher, Waltham, Massachusetts, USA) were used. Reverse transcription and second-strand synthesis were also performed as previously described using the SISPA (sequence-independent single-primer amplification) method [59]. Random amplification of dsDNA was performed using HotStarTaq DNA polymerase (5U/μl; Qiagen, Hombrechtikon, Switzerland) and a 20-cycle protocol to increase the probability of detecting low-frequency viral sequences in samples from passerine birds. After amplification, the samples were purified using the QIAquick PCR Purification Kit (Qiagen, Hombrechtikon, Switzerland) following the manufacturer’s instructions. Finally, the concentration of dsDNA was measured using the Qubit Fluorometer 4 (Invitrogen, Carlsbad, California, USA) [59,60].

### Ticks metagenomic sample processing

#### Tick collection and identification

Twenty-two ticks (n = 19 engorged, n = 3 non-engorged) in adult or nymphal stages were collected from 14 birds belonging to 10 species in the nature reserve Bolle di Magadino and Piano di Magadino, canton Ticino (TI, Switzerland, Tab 2). Ticks were stored at −80°C until further processing. Morphological classification to the species level was achieved via stereomicroscope, following two identification guides provided by Estrada Peña [61,62].

**Table 2:**
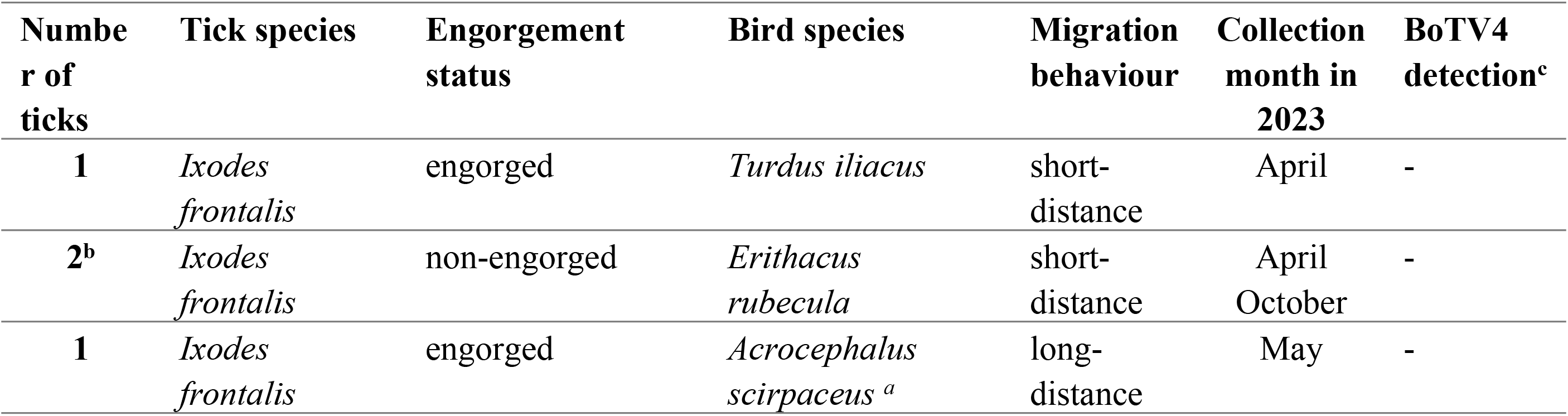
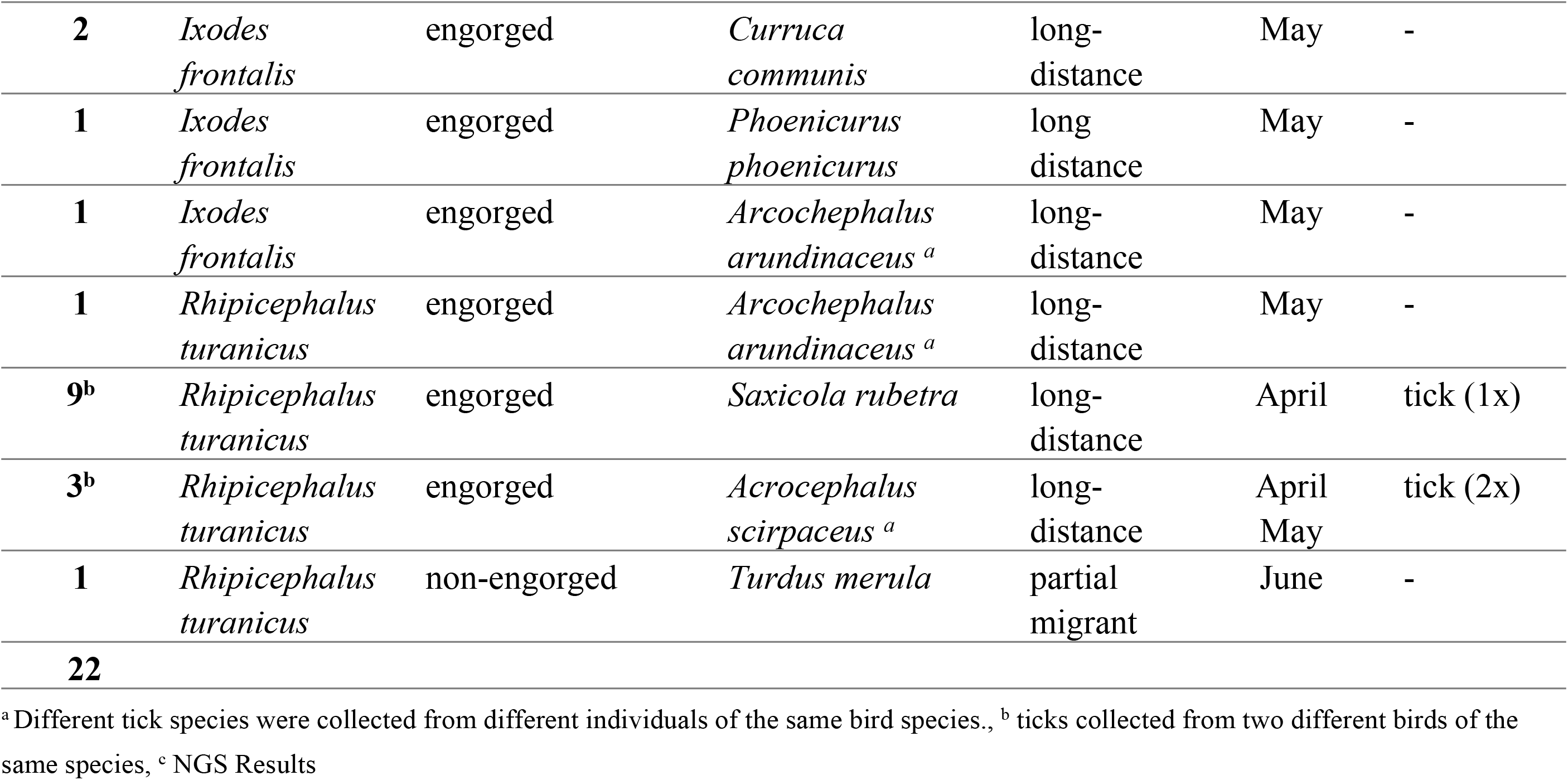
Overview of sampled ticks, their species, feeding status and collection timepoint, the host bird species with corresponding migratory behaviour, and Bole tick virus 4 detection.

| **Number of ticks** | **Tick species** | **Engorgement status** | **Bird species** | **Migration behaviour** | **Collection month in 2023** | **BoTV4 detection<sup>c</sup>** |
|---|---|---|---|---|---|---|
| **1** | *Ixodes frontalis* | engorged | *Turdus iliacus* | short-distance | April | - |
| **2<sup>b</sup>** | *Ixodes frontalis* | non-engorged | *Erithacus rubecula* | short-distance | April<br>October | - |
| **1** | *Ixodes frontalis* | engorged | *Acrocephalus scirpaceus* <sup>*a*</sup> | long-distance | May | - |
| **2** | *Ixodes frontalis* | engorged | *Curruca communis* | long-distance | May | - |
| **1** | *Ixodes frontalis* | engorged | *Phoenicurus phoenicurus* | long distance | May | - |
| **1** | *Ixodes frontalis* | engorged | *Arcochephalus arundinaceus* <sup>*a*</sup> | long-distance | May | - |
| **1** | *Rhipicephalus turanicus* | engorged | *Arcochephalus arundinaceus* <sup>*a*</sup> | long-distance | May | - |
| **9<sup>b</sup>** | *Rhipicephalus turanicus* | engorged | *Saxicola rubetra* | long-distance | April | tick (1x) |
| **3<sup>b</sup>** | *Rhipicephalus turanicus* | engorged | *Acrocephalus scirpaceus* <sup>*a*</sup> | long-distance | April<br>May | tick (2x) |
| **1** | *Rhipicephalus turanicus* | non-engorged | *Turdus merula* | partial migrant | June | - |
| **22** |  |  |  |  |  |  |

<sup>a</sup>Different tick species were collected from different individuals of the same bird species., <sup>b</sup> ticks collected from two different birds of the same species, <sup>c</sup> NGS Results

#### Tick sample processing

After the morphological identification, individual ticks were prepared for NGS. For surface decontamination, each tick was first washed twice with 80% ethanol, dried with a paper towel and then homogenized in 650 μl of Dulbecco’s phosphate-buffered saline (DPBS, St. Louis, Missouri, USA) with a stainless-steel bead (5 mm, Qiagen, Hilden, Germany) using a TissueLyzer II (Qiagen, Hilden, Germany) at 30 Hz for 5 minutes. Afterwards, the resulting homogenate was centrifuged at 16,060 × g for 6 minutes [63].

#### Nucleic acid isolation and SISPA amplification

The 230 μl supernatant was first enriched for virus particles as described for the bird samples. Total nucleic acids (RNA and DNA) were extracted using the QIAamp Viral RNA Mini Kit (Qiagen, Hilden, Germany) with the same modification as previously. Furthermore, tick samples were processed following the same procedures as those applied to the avian samples. The final dsDNA concentration was quantified using the Qubit Fluorometer 4 (Invitrogen, Carlsbad, California, USA) [59,60,63].

#### Library preparation, quality control and sequencing

Downstream processing was conducted at the Next Generation Sequencing (NGS) Platform, University of Bern (Bern, Switzerland). Genomic libraries were constructed using the NEBNext Ultra II DNA Library Prep Kit for Illumina (New England Biolabs, Ipswich, Massachusetts, USA). PhiX174 (PhiX Control v3 Library) was used as a run control according to the manufacturer’s manual (Illumina, San Diego, California, USA). Sequencing was performed on the NovaSeq 6000 system using a high-output flow cell, generating 2 × 150 nucleotide paired-end reads [64].

#### Data analysis

Raw reads were mapped to an in-house database (downloaded in April 2025) containing 73,697 viral sequences from the NCBI nucleotide database using the metagenomic pipeline of SeqMan NGen v. 18 and SeqMan Pro v. 18 software (both DNAStar, Lasergene, Madison, Wisconsin, USA) [64–66]. Only viral species with more than 10 mapped reads per reference genome were considered for downstream analyses. For taxonomic annotation, consensus sequences were compared against the NCBI nucleotide database using BLASTN (v2.17.0) via the NCBI web interface with default parameters, accessed between April and July 2025 [67–69]. Hits were ranked by bit score, retaining only those with an e-value ≤ 1 × 10^−5^ and bit score ≥ 100. Following manual inspection of alignments, specific taxa were assigned based on the top alignment hit (highest bit score). For phylogenetic reconstruction, viral sequences were aligned using MAFFT (v7.526) under default settings (FFT-NS-2 algorithm) [70,71]. Poorly aligned, highly gapping regions were trimmed using trimAl (v1.4.1) with a gap threshold of 90% (-gt 0.9) [72]. Maximum Likelihood (ML) phylogenetic trees were constructed in IQ-TREE 2 (v2.3.4), utilizing the integrated ModelFinder Plus (MFP) algorithm to automatically select the optimal evolutionary substitution model based on the Bayesian Information Criterion (BIC) [73,74]. Branch support was assessed using 1,000 Ultrafast Bootstrap (UFBoot2) replicates, and trees were visualized using the TreeExplorer in MEGA (v12.1) [75,76]. To accurately quantify viral abundances and resolve ambiguous alignments, a non-redundant reference database was constructed by extracting the filtered viral sequences identified via SeqMan NGen (following control subtraction, including phiX174 and Mengo virus) from the previously used NCBI-derived database. Read re-mapping and abundance quantification were performed using RSEM (v1.3.1) with Bowtie2 (v2.5.2) as the alignment backend to allocate multi-mapping reads via an Expectation-Maximization (EM) algorithm [77,78]. Most samples were processed using Bowtie2’s --fast preset with up to 100 reported alignments per read (-k 100) to optimize computational runtime. An initial subset of samples (n=27) was processed using the --sensitive preset with -k 200. The two parameter sets were not systematically compared. Alignment output was converted to BAM format using SAMtools (v1.19.2) [79]. Reads from multiple sequencing lanes per individual bird were summed prior to quantification. For putatively segmented viral genomes, identified heuristically via header patterns (e.g., "segment", "RNA1/RNA2", "L/M/S"), read counts and effective lengths were merged across segments, and abundances were expressed as Reads Per Kilobase per Million mapped reads (RPKM) [80]. The completeness of segment sets (e.g., exactly one L, M and S segment) was not verified for individual viruses. RSEM-derived RPKM values were used for virus detections with RPKM > 0. For the remaining detections, where RSEM returned an RPKM of 0, RPKM was calculated manually from raw read counts, target template length (obtained from the original SeqMan NGen assembly), and total sample depth. Both approaches are conceptually comparable in their normalization logic. All downstream statistical analyses were conducted in R (v4.5.2) [81]. Read abundances (RPKM) were aggregated at the viral family level by summation across assigned taxa per sample. Differences in family-level RPKM abundance between migratory strategies (long-distance vs. short-distance) were evaluated using non-parametric Wilcoxon rank-sum tests with samples (sequencing batches, each comprising one or more birds, with at least one assigned virus detection) as the unit of analysis [82]. Differences in prevalence (proportion of positive samples) were assessed using Fisher’s exact tests [83]. To control the false discovery rate (FDR) arising from multiple hypothesis testing, *p*-values were adjusted using the Benjamini–Hochberg (BH) procedure [84]. Viral families with an adjusted *p*-value (*q*-value) < 0.05 and an absolute log_2_ fold change > 1 were considered statistically significant.

#### Accession Numbers

The raw sequence reads generated in this study are available in the NCBI Sequence Read Archive (SRA) under BioProject accession number PRJNA1459659. Nucleotide sequences of selected viruses identified in this study have been deposited in GenBank under the accession numbers: PX613657 to PX613659 and PX695504 to PX695485.

## Results

A total of 496 birds from 42 species were sampled. Of these, 486 individuals were captured using a catch-and-release method. An additional 10 birds (9 *Corvus corone cornix* and 1 *Falco tinnunculus*) were obtained from various locations across Ticino through population containment plans. From the total cohort, 496 oropharyngeal-cloacal swabs, 164 serum samples, and 22 ticks directly removed from the birds were collected (Tab 1). They belonged to 16 families and five orders, with the majority belonging to the order Passeriformes (90.5%, n=38, S1 Table) [1].

### Avian species diversity and migratory pattern

The captured bird species exhibited diverse migratory behaviour and were initially divided into five categories: resident birds (n=3), partial migrants (n=9), short-distance migrants (n=10), intermediate migrants (n=1), and long-distance migrants (n=19). While capturing the birds, temporal variation in species composition was observed. In the initial phase (April), captures were dominated by short-distance migrants, whereas long-distance migrants were caught in the second phase (end of April to May). Resident species were consistently represented throughout the sampling period. To ensure statistically robust comparisons of migration strategies, only the 19 bird species exhibiting strictly long-distance behaviour were retained in this category. The remaining categories were grouped into resident and short-distance migrants. This resulted in two categories: residents and short-distance migrants (23 species, n=268 samples analyzed, Fig 1A), and long-distance migrants (19 species, n=208 samples analyzed, Fig 1B). Sample numbers ranged widely among species, with *Sylvia atricapilla* (short-distance, n=143) and *Erithacus rubecula* (short-distance, n=53) representing the most frequently sampled species.

**Fig 1:**
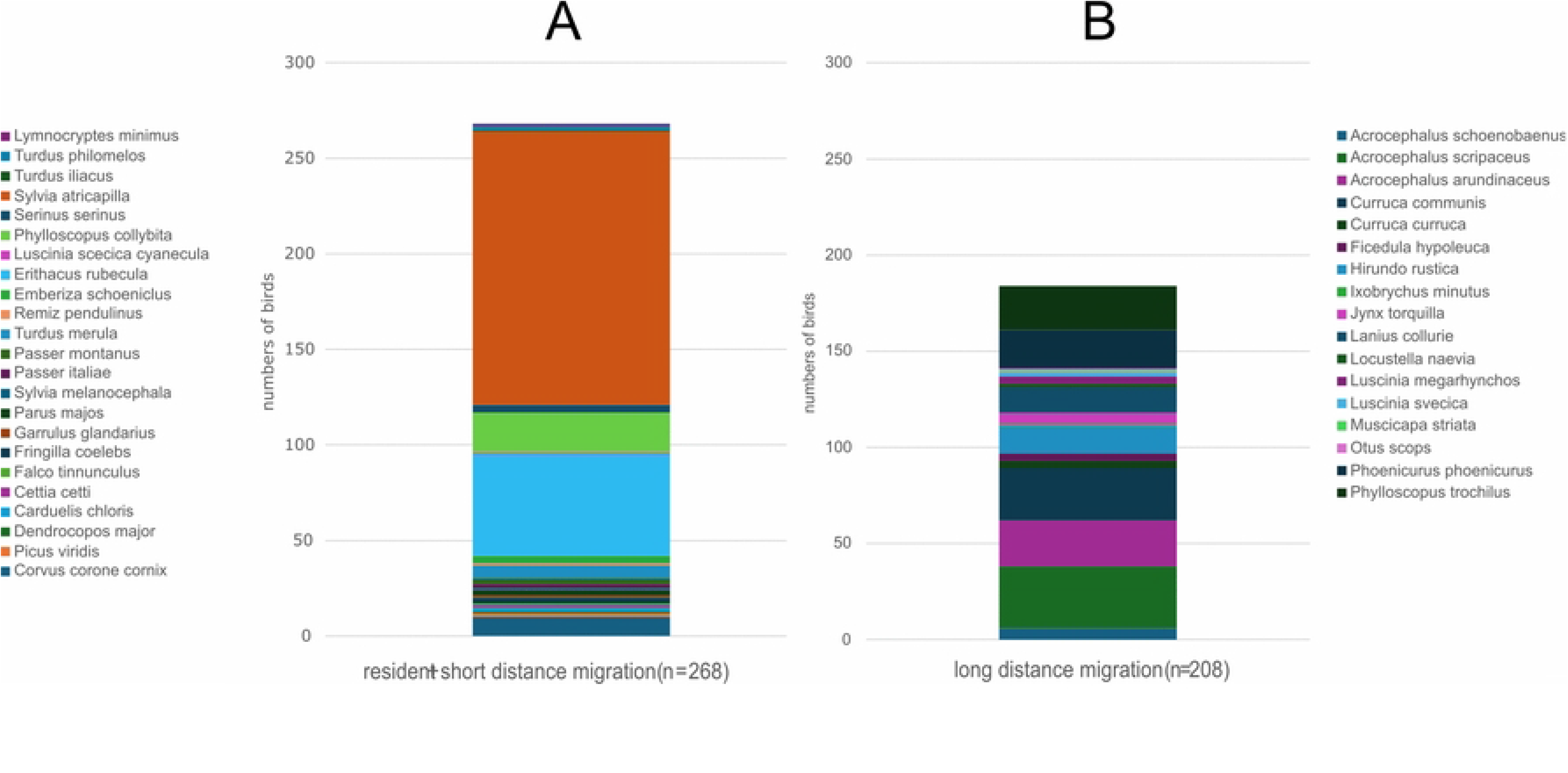
Overview of swab samples distribution across bird migratory strategies. **A**. Stacked column chart shows the number of swab samples (one per individual bird) collected from bird species with short-distance migratory strategies, including resident species and *Lymnocryptes minimus*, a species that may also exhibit long-distance migration. **B**. Stacked column chart shows the number of swab samples (one per individual bird) collected from long-distance migratory species.

### Targeted molecular surveillance of AIV and WNV

First, oropharyngeal-cloacal swabs had been screened using RT-qPCR for two zoonotic pathogens, i.e., Avian Influenza virus (AIV) and West Nile virus (WNV), all of which yielded negative results. Additionally, 164 sera were tested for WNV antibodies using ELISA, yielding 12 doubtful and 15 positive results. To confirm these findings, a WNV serum neutralization test (SNT) was performed, starting at 1:32 serum dilution, due to the limited amount of available material. However, no neutralizing antibodies were detected (S2 Table). Further specific SNTs for other Orthoflaviviruses were not possible due to the limited sample quantity.

### Virome Composition and Host-Specific Distribution

Metagenomic sequencing of 476 oropharyngeal-cloacal swab samples generated 3.56 × 10^9^ raw reads, with individual pools yielding between 7 × 10^6^ and 4 × 10^7^ reads. Bioinformatics analysis identified 6.22 × 10^5^ viral reads. After filtering for quality (E-value ≤10^−5^, bit score ≥ 100) and robustness (> 10 mapped reads per reference), 4.32 × 10^5^ classified viral reads remained for analysis. These were assigned to 17 RNA (58.6%) and 12 DNA families (41.4%), encompassing a diverse virome of both single-stranded (ss) and double-stranded (ds) genomes. RNA viruses exhibited higher family level diversity but were less abundant per individual, whereas DNA viruses were less diverse yet more prevalent in read counts and across bird species. Overall, viral reads represented only a minor fraction of the total sequencing output. The viral reads were assigned to 29 distinct virus families and a group of unclassified viruses, which was further divided into three subgroups, i.e. *Cressdnaviricota*, other DNA viruses, and RNA viruses not specifically classified in the NCBI Taxonomy Browser [85] (Fig 2). Among the 42 bird species, *S. atricapilla* (n = 143) had both the highest number of collected samples and the highest viral diversity, with 17 families. Interestingly, viral richness did not strictly correlate with sample size, *E. rubecula* (n = 53) hosted only nine virus families despite its large representation. By contrast, *Hirundo rustica* (long-distance, n=14) hosted 11 families, while *Turdus merula* (partial migrant, n=7) and *Jynx torquilla* (long-distance, n=6) each harboured 10 families. The lowest viral richness was observed in *Muscicapa striata* (long-distance, n=1), with viral reads assembled only to the family *Circoviridae*.

**Fig 2.**
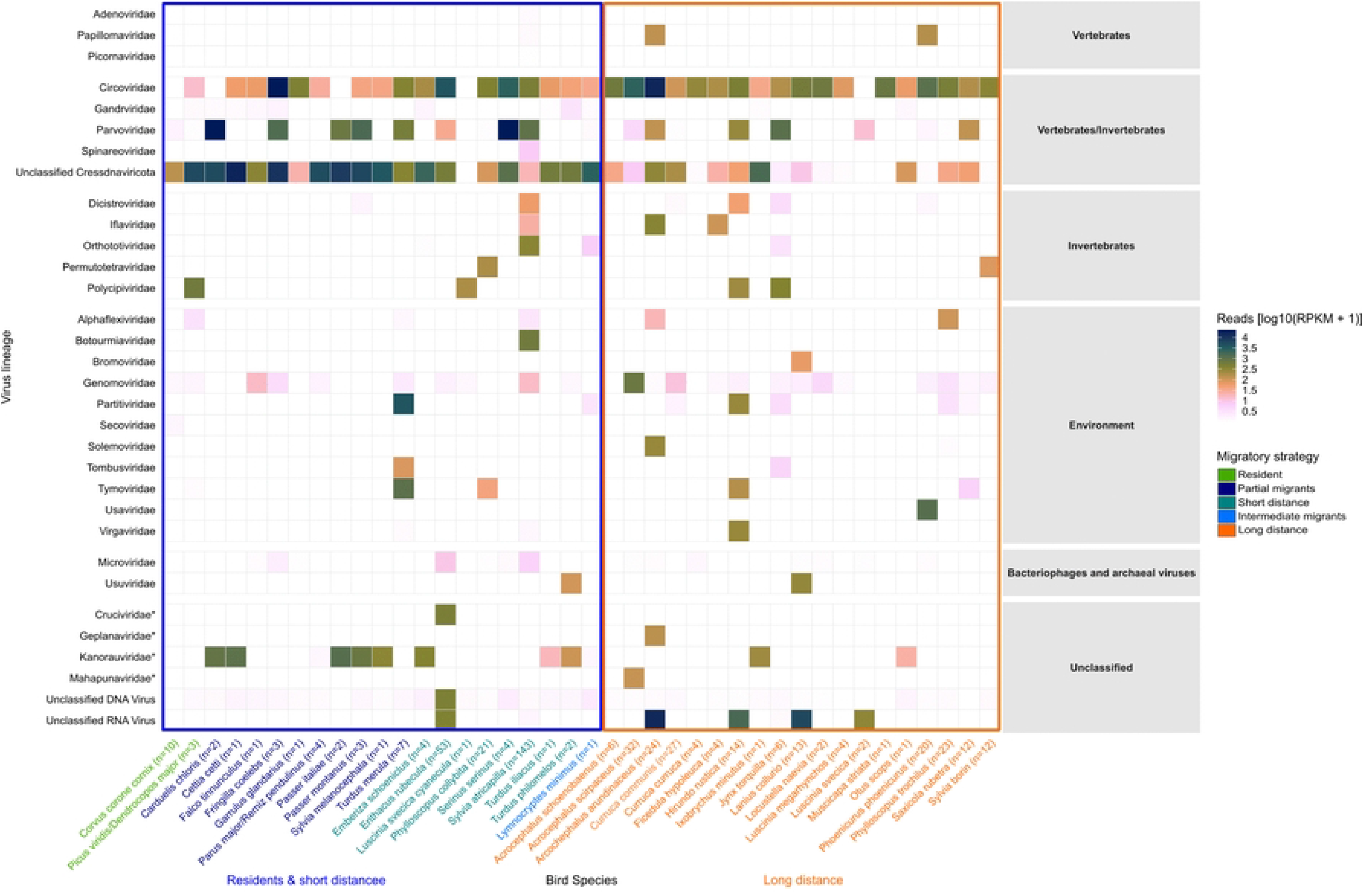
Distribution of the virome across the sampled bird species. The heatmap displays 29 viral families and three groups of unclassified viruses across 42 avian species (three resident and 39 migratory species), representing five migratory strategies ranging from resident to long-distance migrants. Viral lineages are displayed on the y-axis and bird species on the x-axis. Colour intensity reflects relative viral abundance [log_10_(RPKM + 1)]. White tiles indicate zero detected reads, whereas increasing colour intensity represents higher viral abundance. * According to the current ICTV taxonomy, these families belong to the phylum *Cressdnaviricota*. Given their phylogenetic relationship and frequent detection in environmental samples, they are thought to be classified as environmental.

### Prevalence and Abundance Profile of Avian Viral Families

While the species-level heatmap (Fig 2) highlights host-specific detection patterns across individual avian taxa, aggregating data at the viral family level provides a global overview of the entire virome structure (Fig 3). For this, the prevalence of each family (percentage of positive samples) was plotted against mean viral abundance (log_10_(RPKM + 1)) across migratory strategies. The analysis was based on samples (sequencing batches) with at least one assigned virus detection (*n*=70 short-distance migrants and *n*=66 long-distance migrants). Single-stranded DNA viral families, specifically *Circoviridae, Genomoviridae, Parvoviridae*, and unclassified *Cressdnaviricota*, represented the highest overall host species prevalence. Among these, *Circoviridae* and *Parvoviridae* also had the highest mean RPKM values, making them, along with unclassified RNA viruses, dominant, highly abundant components of the wild bird virome. Except for these dominant groups, other vertebrate-infecting viral families generally exhibited lower detection frequencies across migratory species. Invertebrate- and plant-associated viruses as well as bacteriophages displayed lower overall prevalence patterns.

**Fig 3.**
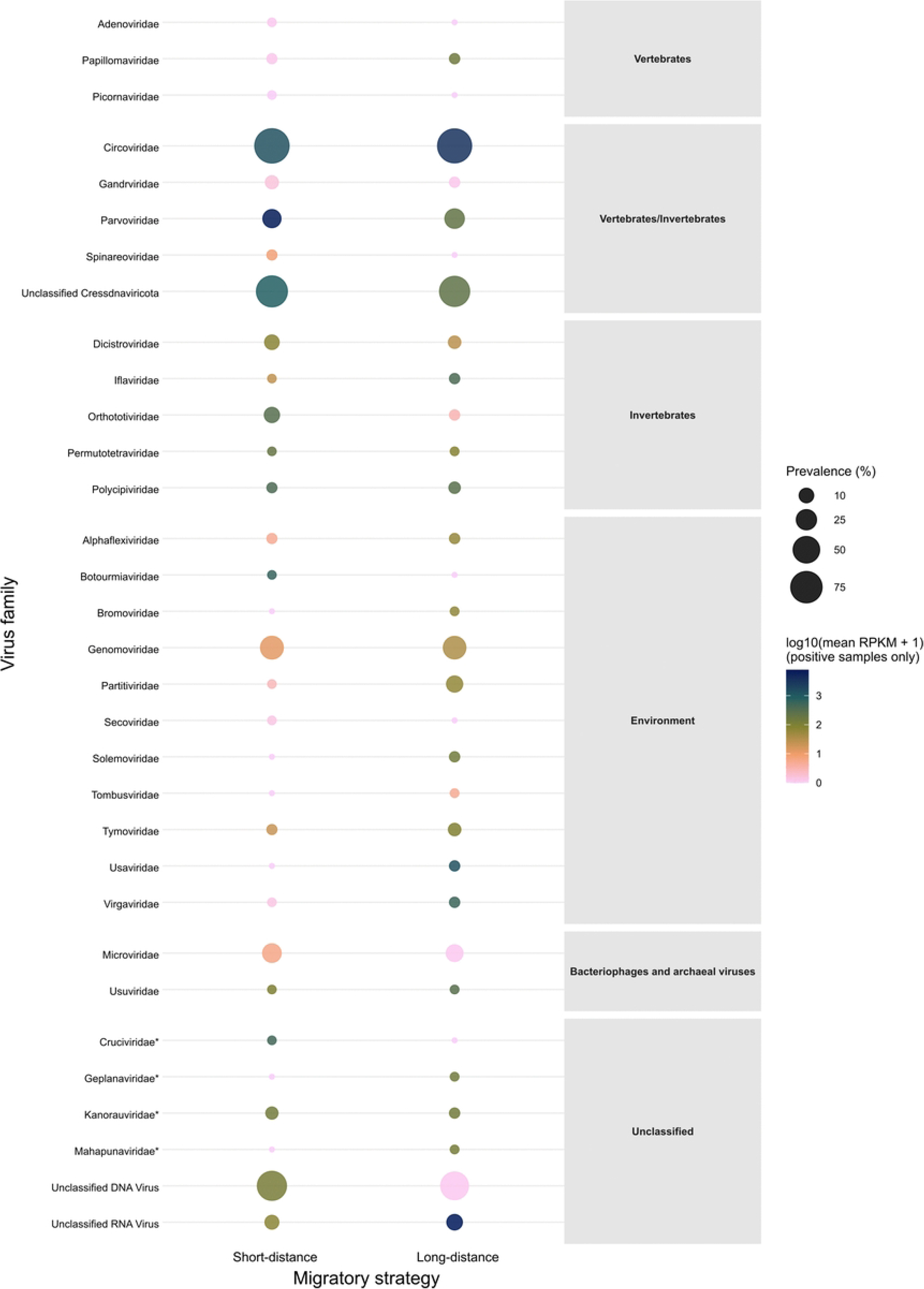
Prevalence and abundance profile of avian viral families. Bubble plot displays, for each viral family, the prevalence (percentage of positive samples) plotted against mean viral abundance (log_10_(RPKM + 1), calculated among positive samples only) across migratory groups (short-distance/resident vs. long-distance migrants). Bubble size corresponds to prevalence, while color intensity reflects mean viral abundance among positive samples. Families are grouped by host category. * According to the current ICTV taxonomy, these families belong to the phylum *Cressdnaviricota*. Given their phylogenetic relationships and frequent detection in environmental samples, they are thought to be classified as environmental. Differences in prevalence between migratory groups were tested using Fisher’s exact test with Benjamini-Hochberg correction. No family differed significantly.

For further analysis, virus families were categorized based on their primary hosts: (i) vertebrates, (ii) invertebrates, (iii) environment, (iv) bacteriophages, and (v) newly established and unclassified families.

### Vertebrate-associated viruses

Seven virus families were identified based on reads assembled to viral genomes known to infect vertebrates, mainly mammals and birds: *Adenoviridae, Circoviridae, Gandrviridae, Papillomaviridae, Parvoviridae*, *Picornaviridae*, and *Spinareoviridae*, alongside unclassified viruses of the phylum *Cressdnaviricota.* This category had the highest viral abundance in the study, with 2.05 × 10^5^ reads. Notably, *Circoviridae* dominated with 1.85 × 10^5^ reads, predominantly in *H. rustica* (9.96 × 10^4^ reads) and *Acrocephalus arundinaceus* (long-distance, 2.73 × 10^4^ reads). Overall, *Circoviridae* and the unclassified *Cressdnaviricota* group (1.05 × 10^4^ reads) were detected in nearly all species (37/42). *Parvoviridae* (9.56 × 10^3^ reads) was detected across 21 bird species, while *Picornaviridae* (11 reads) was the least abundant, detected only in *S. atricapilla*.

### Invertebrate-associated viruses

Reflecting the diet of most investigated species (40/42), five RNA viral families primarily infecting insects were identified: *Dicistroviridae, Iflaviridae, Orthototiviridae, Permutotetraviridae*, and *Polycipiviridae.* The overall read count was 1.31 × 10^4^, with high concentrations in specific species, particularly in *S. atricapilla* (2.9 × 10^3^ reads for *Dicistroviridae*) and *J. torquilla* (6.96 × 10^3^ reads for *Polycipiviridae*). In contrast, *Permutotetraviridae* (60 reads) showed the lowest prevalence, detected only in *Phylloscopus collybita* (short-distance) and *Sylvia borin* (long-distance). Additionally, some *Parvoviridae* reads likely correspond to the subfamily *Densovirinae*, which specifically infect arthropod hosts were also detected.

### Environment-associated viruses

The category with the highest family richness comprised 11 families predominantly associated with plant and fungal hosts: *Alphaflexiviridae*, *Botourmiaviridae*, *Bromoviridae*, Genomoviridae, *Orthototiviridae*, *Partitiviridae*, *Secoviridae*, *Solemoviridae*, *Tombusviridae*, *Tymoviridae*, and *Virgaviridae*. Although this group represented 39.3% of the total family richness (11 out of 28 identified families), its cumulative abundance was relatively low, accounting for only 6.4% of the total classified viral reads (2.78 × 10^4^ reads). Within this category, five families were identified in only a single bird species. However, *Genomoviridae* (1.81 × 10^4^) was identified in more than half of the species (23/42), with its highest viral load in *S. atricapilla* (1.37 × 10^4^ reads) and *Curruca communis* (long-distance, 1.7 × 10^3^ reads).

### Bacteriophages and archaeal viruses

Although this study targeted eukaryotic viruses and included nuclease treatment to reduce bacterial and host nucleic acids, sequences mapping to bacteriophages and archaeal viruses were also recovered. Specifically, sequences belonging to the phage family *Microviridae* (515 reads) were identified in 14 of the 42 bird species. Furthermore, reads mapping to the *Usuviridae* family (643 reads), which specifically infect methanogenic gut archaea, were detected in two bird species.

### Newly established and unclassified families

This last category included recently established taxa within the phylum *Cressdnaviricota* such as C*ruciviridae* (11 reads, 1/42 species), *Mahapunaviridae* (100 reads, 1/42 species), and *Kanorauviridae* (212 reads, 12/42 species), and *Geplanaviridae* (26 reads, 1/42 species) [86,87]. The unclassified DNA viruses represented the highest overall sequence abundance (3.71 × 10^5^ reads, 35/42 species), dominated by *E. rubecula* with 3.59 × 10^5^ reads, and *Corvus corone cornix* (resident, 1.04 × 10^4^ reads). In contrast, unclassified RNA viruses (4.97 × 10^3^ reads) were less prevalent, detected in only seven species.

### Differential Viral Abundance Across Migratory Strategies

To determine whether specific viral families were differentially abundant between short-distance and long-distance migrants, two-sided Wilcoxon rank-sum tests with FDR correction were performed (Fig 4A). All detected viral families clustered below the significance threshold (FDR < 0.05, |log_2_FC| >), indicating no statistically significant difference in viral family abundance between migratory strategies. Furthermore, while both DNA and RNA viral families were broadly distributed across positive and negative fold changes, no overall bias linked to viral genome architecture was observed. Additionally, the comparison of eight key viral families regarding viral abundance (log_10_(RPKM + 1)) across short- and long-distance migrants confirmed the absence of statistically significant differences across migratory strategies (Fig 4B). *Circoviridae* and *Parvoviridae* exhibited high viral loads across both migratory groups, mirroring the high overall prevalence patterns observed previously (Fig 3).

**Fig 4.**
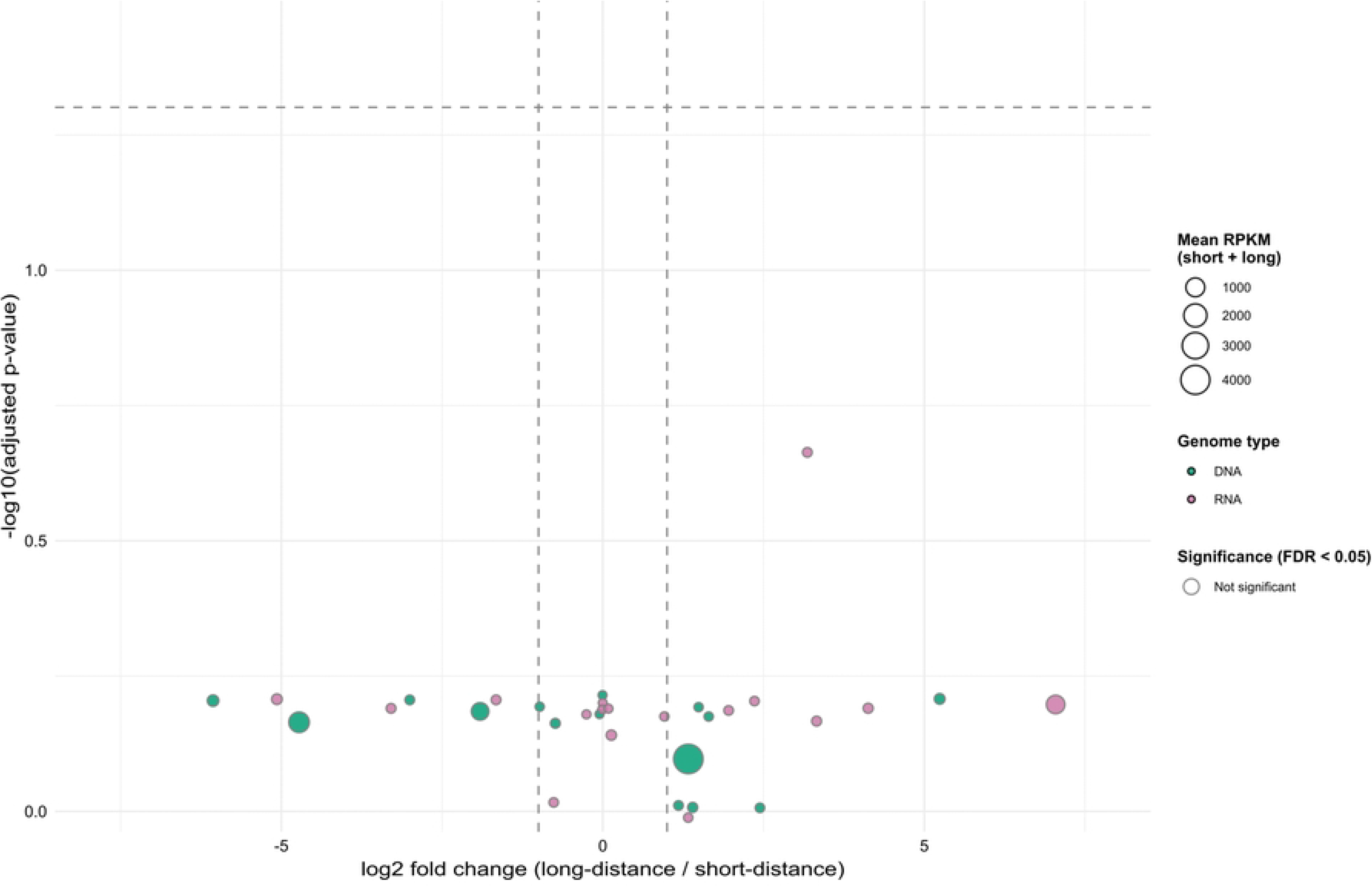
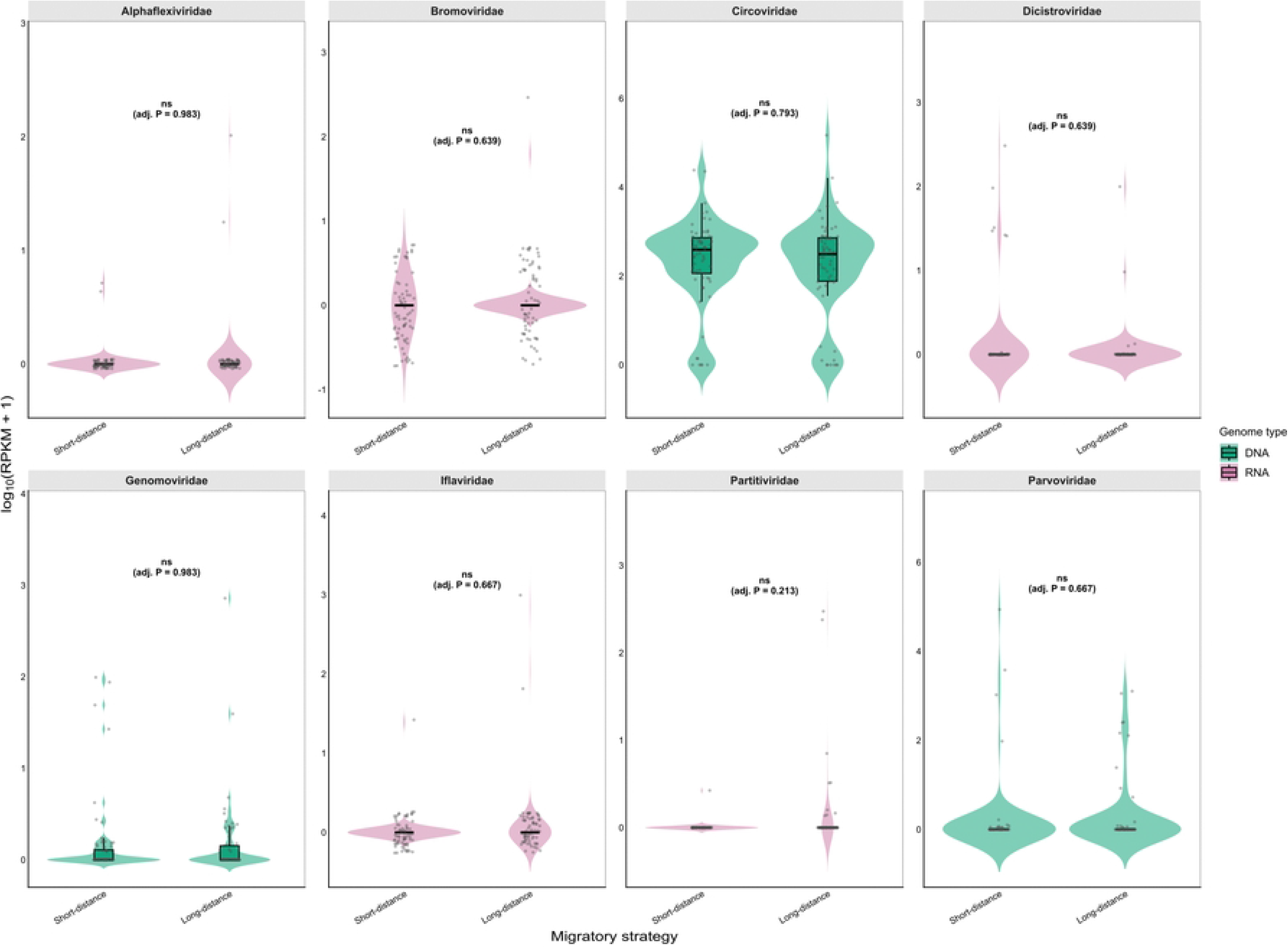
Abundance and differential analysis of viral families across migratory strategies. Statistical significance for each viral family was assessed using two-sided Wilcoxon rank-sum tests across samples, with *p*-values adjusted for multiple testing using the Benjamini–Hochberg procedure to control the false discovery rate (FDR). **A.** Volcano plot illustrating differential abundance of viral families between long-distance and short-distance migrants. The x-axis shows the log_2_ fold change in mean RPKM abundance (long-distance vs. short-distance migrants) and the y-axis shows statistical significance (−log_10_ adjusted *p*-value). Points are filled by genome type (DNA in green, RNA in pink) and outlined by significance category (orange: higher in long-distance migrants, blue: higher in short-distance migrants, grey: not significant). No family reached significance. Point size represents the summed mean RPKM abundance across both migratory groups. Dashed lines indicate a *p*-value of 0.05 (-log_10_=1.3), and the vertical lines mark a log_2_ fold change of -1 and +1 (two-fold difference). Families were considered significant only if they fulfilled both criteria. **B.** Violin plots displaying the abundance (log_10_(RPKM + 1)) of eight key viral families in short-distance vs. long-distance migrants. Violin shapes represent the density distribution across samples, overlaid with boxplots; the central horizontal line indicates the median, and the box spans the interquartile range (25^th^–75^th^ percentile). Whiskers extend to the most extreme values within 1.5 × the interquartile range from the box. Values beyond this range remain visible as individual jittered data points. Each point corresponds to one sample (sequencing batch). Numbers above each panel show the adjusted p-value (ns, not significant). Panels are color-coded by genome type (DNA in green, RNA in pink).

### Diversity and phylogeny of avian DNA viruses

To further investigate the evolutionary relationships of the identified DNA virome, representative viral sequences with sufficient genome coverage from the most prevalent families were analyzed (Fig 5 and 6).

**Fig 5.**
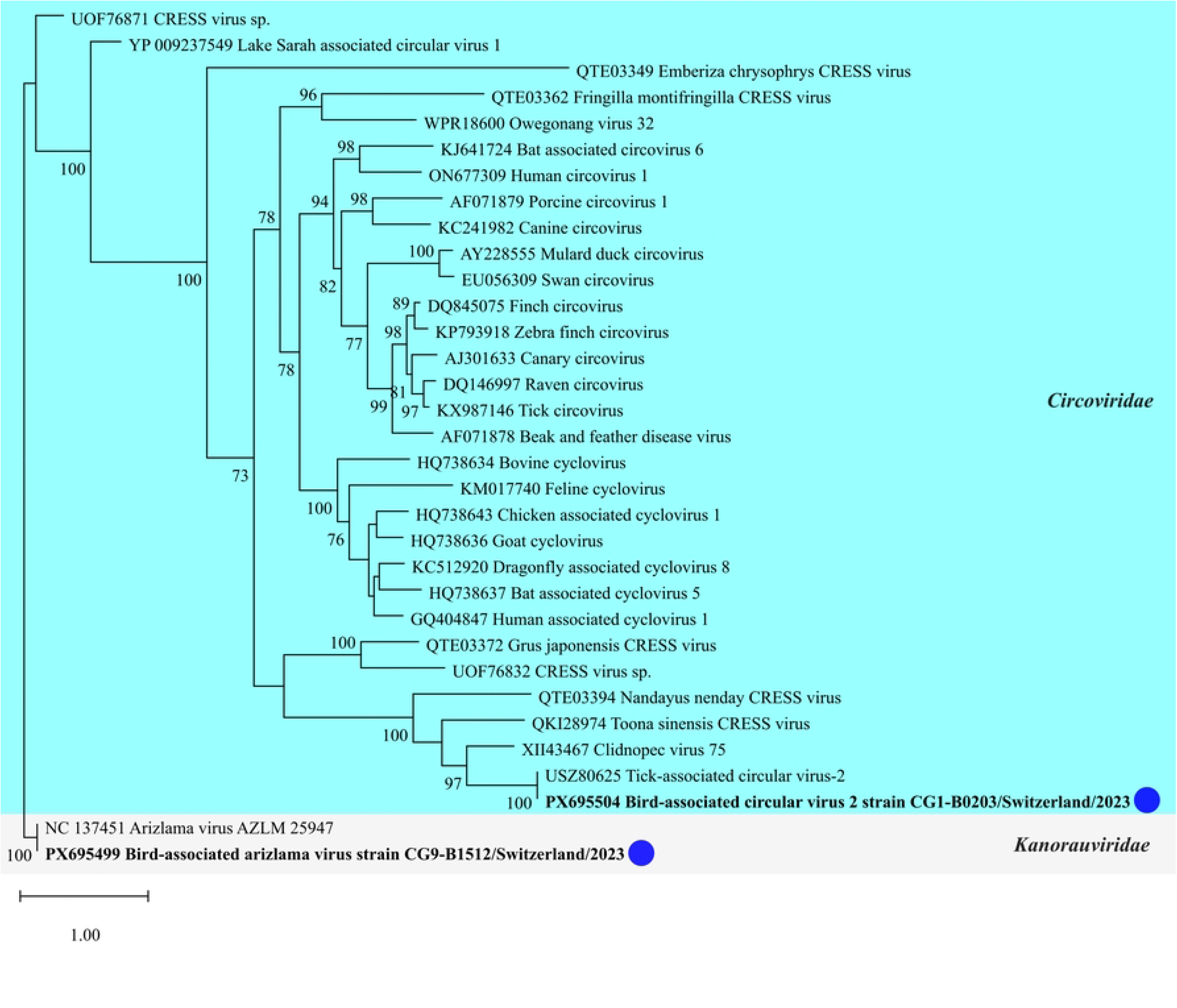
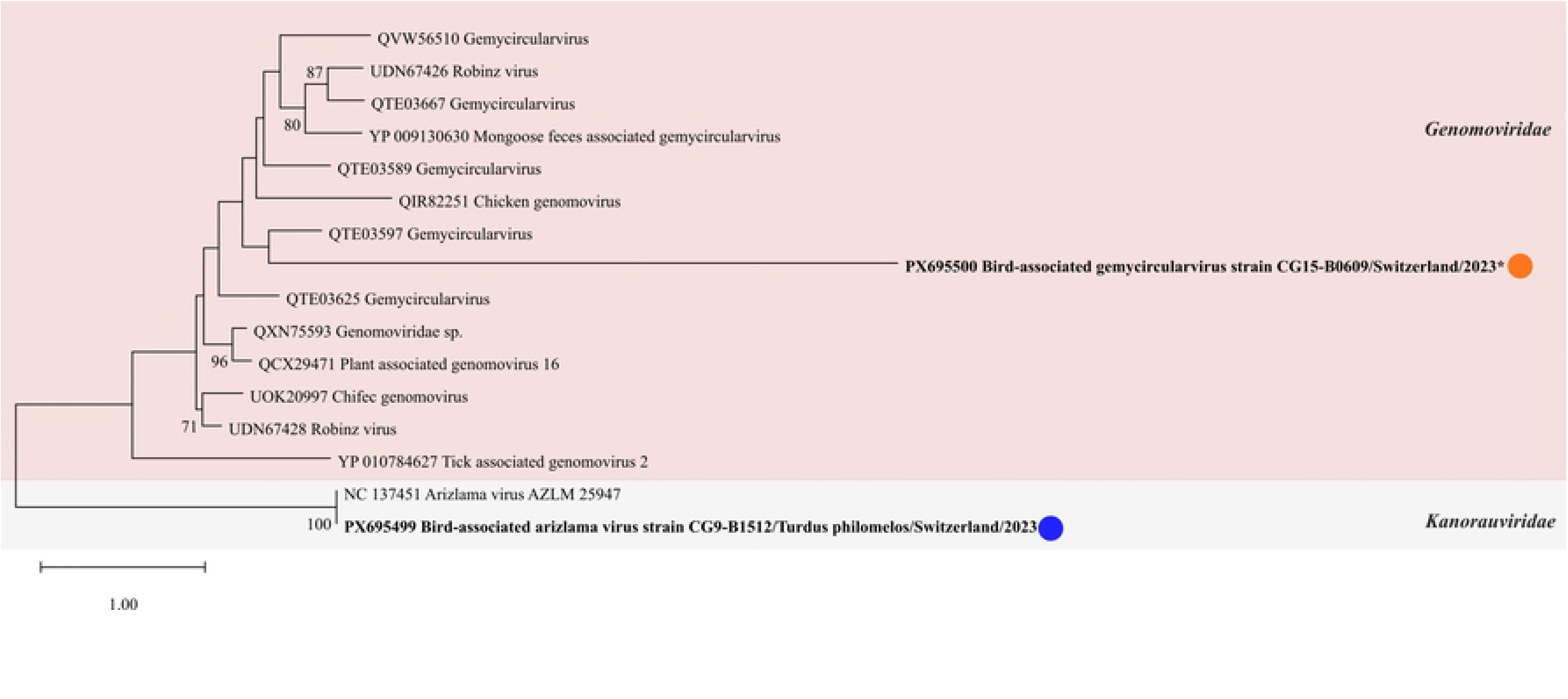
Phylogenetic analysis of the phylum *Cressdnaviricota*. The Maximum Likelihood (ML) tree was constructed using IQ-TREE 2 (v2.3.4) based on an alignment of amino acid (aa) sequences of the replication-associated protein (Rep) performed using MAFFT (v7.526), followed by alignment trimming with trimAl (gap threshold: 0.9) to retain highly conserved domains. Numbers at the internal nodes represent bootstrap support values derived from 1,000 Ultrafast Bootstrap (UFBoot2) replicates. For clarity, only bootstrap values ≥ 70% are displayed. Sequences identified in this study are shown in bold and annotated with a blue (short-distance migrant) or orange (long-distance migrant) dot. Asterisks (*) indicate partial Rep sequences. Both trees are rooted using Arizlama virus (NC_137451) as an outgroup. **A**. Representative members of the families *Circoviridae* (including the genera *Circovirus* and *Cyclovirus*) and *Kanorauviridae* were included for taxonomic context. **B**. Representative reference sequences for the families *Genomoviridae* and *Kanorauviridae* were included for taxonomic context.

**Fig 6.**
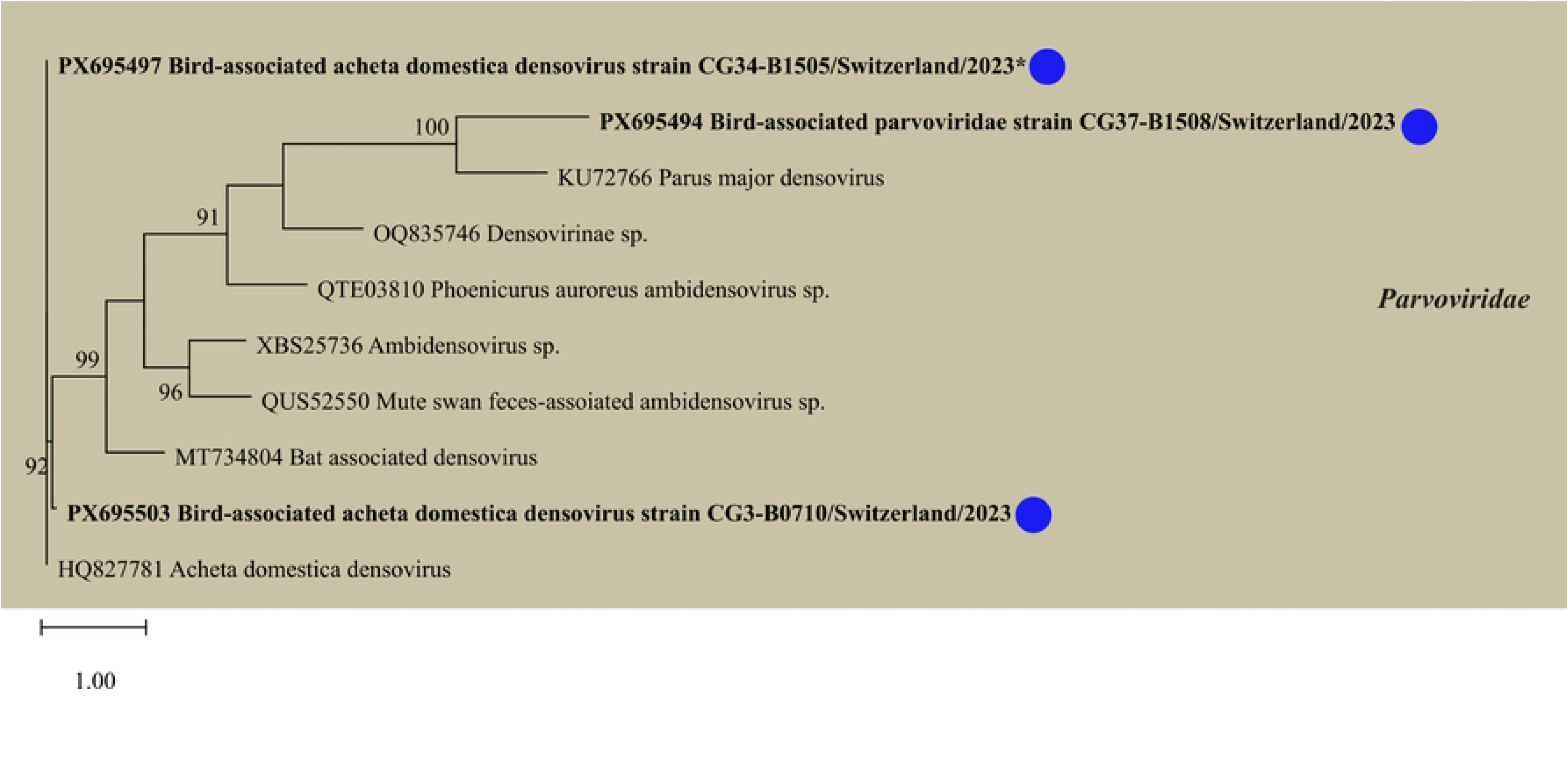
Phylogenetic analysis of *Parvoviridae*. The Maximum Likelihood (ML) tree was constructed using IQ-TREE 2 (v2.3.4) based on an alignment of amino acid (aa) sequences of the non-structural protein (NS1) performed using MAFFT (v7.526), followed by alignment trimming with trimAl (gap threshold: 0.9) to retain highly conserved domains. Numbers at the internal nodes represent Ultrafast Bootstrap (UFBoot2) support values derived from 1,000 replicates. For clarity, only bootstrap values ≥ 70% are displayed. Representative members of the subfamily *Densovirinae* belonging to the family *Parvoviridae* were included for taxonomic context. Sequences identified in this study are shown in bold and annotated with a blue (short-distance migrant) dot. Asterisks (*) indicate partial NS1 sequences.

### Phylum Cressdnaviricota

Viruses belonging to phylum *Cressdnaviricota* (circular replicase-encoding single-stranded DNA viruses) represented a prominent and highly diverse component of the study’s findings. Within the family Circoviridae, viral sequences were widely distributed and detected in 37 of 42 bird species. Most identified consensus sequences were closely related to the tick-associated circular virus 2. Notably, a complete genome sequence (<u>PX695504</u>; 2,123 nt) recovered from *S. atricapilla* showed 99.7% identity to a previously identified tick-associated circular virus 2 strain (<u>ON456885</u>) and features the typical two open reading frame (ORF) structure encoding the capsid and Rep proteins. Additionally, to the classical circoviruses, a highly divergent partial sequence (<u>PX695499</u>; 795 nt) was identified in *A. arundinaceus* with 97.5% identity to an Arizlama virus (<u>NC_137451</u>) and contained a single ORF encoding replication-associated (Rep) protein. This sequence is classified within the family *Kanorauviridae*, bypassing the classical family boundaries (Fig 5A). Members of the family *Genomoviridae* were also identified in several short- and long-distance migratory bird species. Specifically, a partial consensus sequence (<u>PX695500</u>; 1,222 nt) had 98% identity to a previously detected *Gemycircularvirus* (<u>MH378453</u>). This sequence contained two partial ORFs encoding capsid (Cap) and Rep protein (Fig 5B).

### Parvoviridae

Within the subfamily *Densovirinae*, a nearly complete consensus genome sequence (<u>PX695503</u>; 3,980 nt) identified in *S. serinus* and a partial sequence (<u>PX695497</u>; 1,649 nt) detected in *P. italiae* showed 99.8% identity to *Acheta domesticus* densovirus (AdDNV, KF015278), a known insect pathogen [88]. The first sequence contains two ORFs encoding NS1 and NS2, along with partial ORFs for NS3, VP1, and VP2 (structural proteins). The second contains an ORF for NS2 and a partial NS1 ORF. A third sequence (<u>PX695498</u>; 865 nt) shared 99% identity with a previously identified *Grus japonensis* parvoviridae sp. (PQ438377) and contained one partial ORF encoding NS1 protein (Fig 6).

### Diversity and phylogeny of avian RNA viruses

To analyze the evolutionary diversity and host relationships of the RNA virome, representative sequences with sufficient genome coverage across key RNA viral families were subjected to phylogenetic reconstruction (Fig 7).

**Fig 7.**
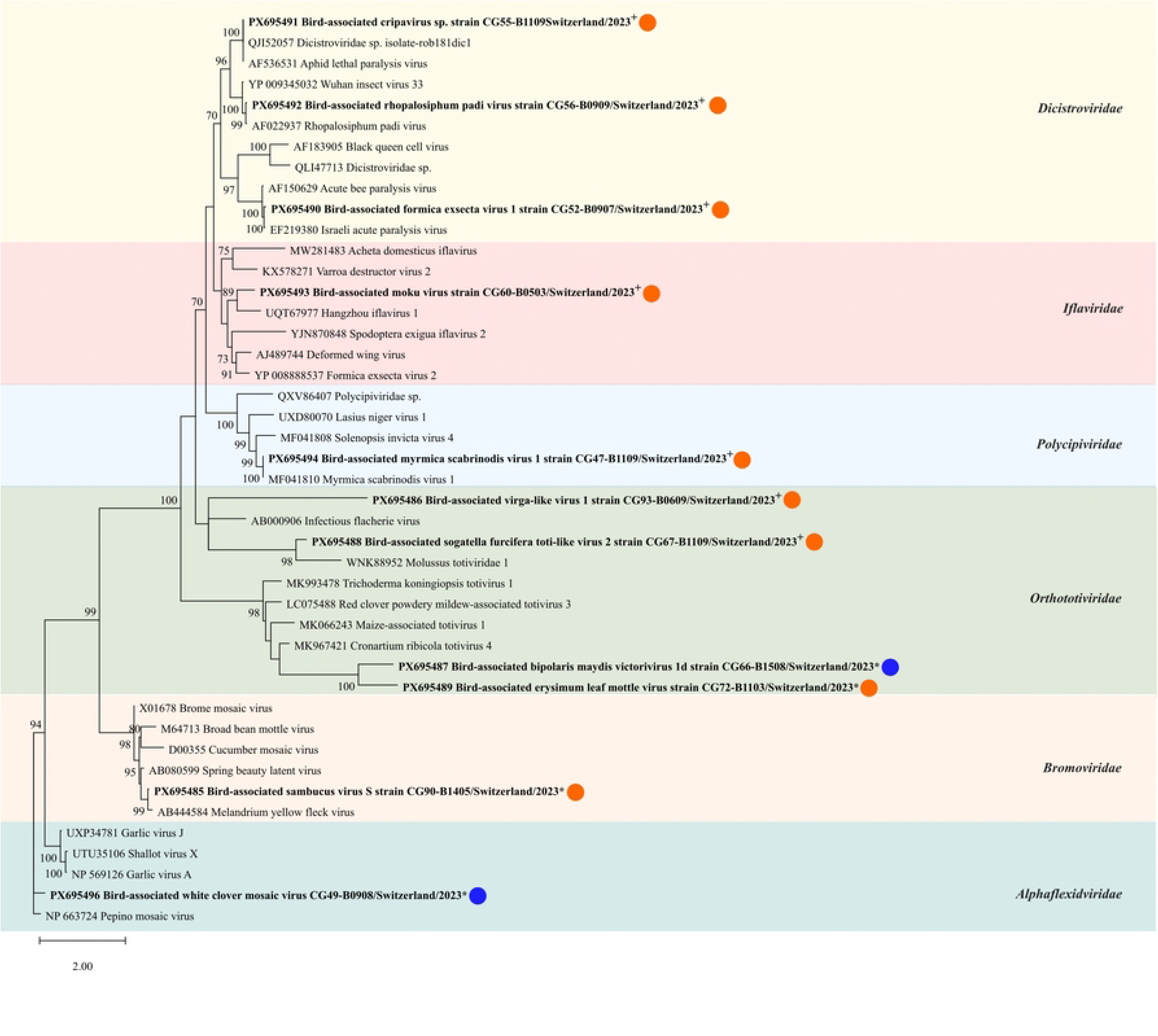
Phylogenetic analysis of RNA genomes. The Maximum Likelihood (ML) tree was constructed using IQ-TREE 2 (v2.3.4) based on an alignment of amino acid (aa) sequences of the RNA-dependent RNA polymerase (RdRp) performed using MAFFT (v7.526), followed by alignment trimming with trimAl to retain highly conserved functional domains. Numbers at the internal nodes represent Ultrafast Bootstrap (UFBoot2) support values derived from 1,000 replicates. For clarity, only bootstrap values ≥ 70% are displayed. Representative members of different RNA virus families were included for taxonomic context. Sequences obtained in this study are indicated in bold and annotated with a blue (short-distance migrant) or orange (long-distance migrant) dot. * represents the environment-infecting viruses, while ^+^ indicates invertebrate-infecting viruses.

### Invertebrate-associated viruses

A total of seven consensus sequences were assembled and classified into four viral families: *Dicistroviridae, Orthototiviridae, Iflaviridae*, and *Polycipiviridae* (Fig 7), with five detected in long-distance migrants. Within the family *Dicistroviridae*, three sequences encoding structural and non-structural proteins were identified. The first identified in *S. atricapilla* (<u>PX695490</u>; 2,209 nt) showed 95% identity to a Formica exsecta virus (<u>NC_023021</u>). The second from *S. atricapilla* (<u>PX695492</u>; 2,144 nt) exhibited 99.66% identity to a *Rhopalosiphum padi* virus (<u>MN535735</u>). Finally, a third sequence detected in *J. torquilla* (<u>PX695491</u>; 2,193 nt) had 96% identity to a Cripavirus (<u>PP872551</u>). In *Orthototiviridae*, two sequences containing partial RdRp ORFs were detected. One identified in *H. rustica* (<u>PX695486</u>; 2,334 nt) showed 96% identity to a known Broome virga-like virus (<u>MT498833</u>), and another identified from *J. torquilla* (<u>PX695488</u>; 1,102 nt) exhibited 89% identity to a *Sogatella furcifera* toti-like virus 2 (<u>ON402806</u>). A nearly complete *Polycipiviridae* genome sequence (<u>PX695494</u>; 11,622 nt) from *J. torquilla* showed 96% identity to a *Myrmica scabrinodis* virus (<u>NC_035457</u>), with ORFs encoding capsid, hypothetical, and RdRp proteins. The final invertebrate-associated host virus was detected in *A. arundinaceus* (<u>PX695493</u>; 1,812 nt) and assigned to *Iflaviridae* with 99% identity to *Vespa velutina* Moku virus (<u>MF346349</u>), covering part of a polyprotein ORF.

### Environment-associated viruses

A total of four consensus sequences were mapped, which were detected in both short-distance and long-distance migrants (Fig 7). Within the family *Alphaflexiviridae,* a sequence (<u>PX695496</u>; 4,006 nt) was identified in *S. atricapilla* and showed 95% identity to White clover mosaic virus (<u>PQ162444</u>). This sequence contained partial ORFs for replication and coat proteins, as well as triple gene block (tgb1, tgb2, tgb3). In *Bromoviridae*, a sequence (<u>PX695485</u>; 831 nt) was assembled from *Lanius collurio* (long-distance), sharing 96% identity with Sambucus virus 2 (<u>NC_040389</u>) and containing a partial ORF for protein 2a. Additionally, two sequences showed strong associations with the recently established family *Orthototiviridae*. In *Turdus merula,* a sequence (<u>PX695489</u>; 1,892 nt) shared 91% identity to *Erysimum leaf* mottle virus (ErLMV, <u>OL420671</u>). Notably, while this virus is taxonomically assigned to the *Tymoviridae*, in our phylogenetic analysis it clustered within the *Orthototiviridae* clade. This sequence encoded partial ORFs for replication and movement protein. Further, in *Lymnocryptes minimus* (intermediate migrant*, n=*1), a sequence (<u>PX695487</u>; 1,082 nt) was found that shared 92% identity with a *Bipolaris maydis* victorivirus (<u>MN935505</u>), including a partial RdRp ORF.

### Phylogenetic investigation of Bole tick virus 4 (BoTV4)

To investigate the potential role of birds as dispersal vectors for tick-borne pathogens, we analyzed 22 ticks (19 engorged) removed from captured birds. In total, 14 birds representing ten species were found to be tick-infested (Tab 2). Seven of these host species exhibited long-distance migratory behaviour, wintering in central or southern Africa, while the three remaining species only migrate to the Mediterranean region and North Africa [33]. Nearly all ticks (21/22) were collected between April and June, closely coinciding with the arrival of birds from Africa. Shotgun metagenomic sequencing of the ticks generated a total of 5.03 × 10^8^ raw reads, of which 7.52 × 10^4^ viral reads mapped to four viral families: *Circoviridae, Pestiviridae*, *Nairoviridae,* and *Partitiviridae*. Notably, no major tick-borne human pathogen sequences including Tick-borne encephalitis virus (TBEV), were detected. However, 99.73% of the viral reads (7.5 × 10^4^) were classified into the family *Pestiviridae*. These reads were detected in three engorged *Rhipicephalus turanicus* ticks. A complete consensus genome sequence (<u>PX613658</u>; 16,230 nt) and a nearly complete sequence (<u>PX613659</u>; 16,181 nt), both sharing 95% nt identity with a Bole tick virus 4 (BoTV4) strain originally identified in China (<u>MW561976</u>), were assembled. These genomes encode a single long ORF for a polyprotein. Additionally, a third partial sequence (<u>PX613657</u>; 1,653 nt) showed 94% nt identity to another strain from China (<u>PP945237</u>). Phylogenetic analysis based on conserved aa domains of the RdRp (Fig 8A) revealed that our sequences form a robust monophyletic cluster with other previously identified BoTV4 strains, officially placing them within the newly established genus *Boletivirus* within the family *Pestiviridae* (order *Amarillovirales*). Importantly, prior to the major taxonomic reorganization in 2025, BoTV4 and related large-genome flavi-like viruses were classified as unclassified members of the family *Flaviviridae*. This was because the genera *Pestivirus, Hepacivirus*, and *Pegivirus* were accommodated as lower-rank taxa within the *Flaviviridae*. The recent taxonomic elevation of these lineages to distinct family levels successfully separates them from the classical, vertebrate-restricted *Flavivirida*e by establishing three monophyletic groups: Clade I (*Flaviviridae*), Clade II (*Pestiviridae*), and Clade III (*Hepaciviridae*) [89]. Reflecting this evolutionary split, our sequences belonging to Clade II (*Pestiviridae*) showed no close phylogenetic relationship to Clade I lineages, such as the classical unsegmented genus *Orthoflavivirus* and the highly divergent genus *Jingmenvirus*, which comprises segmented tick-borne viruses such as Alongshan virus (ALSV) [90,91]. Crucially, our phylogenetic trees demonstrate that the *Boletivirus* (BoTV4) cluster is highly divergent from all other established genera within the Pestiviridae family (such as *Orthopestivirus*), forming a distinct evolutionary lineage (Fig 8A and 8B). Detailed analysis reinforces the global clustering of Swiss BoTV4 consensus sequences with BoTV4 strains from China, Russia, and Africa. While the group forms a robust monophyletic cluster (bootstrap support of 96), the short internal branch lengths indicate a high degree of sequence conservation across these distant geographic regions (Fig 8B).

**Fig 8.**
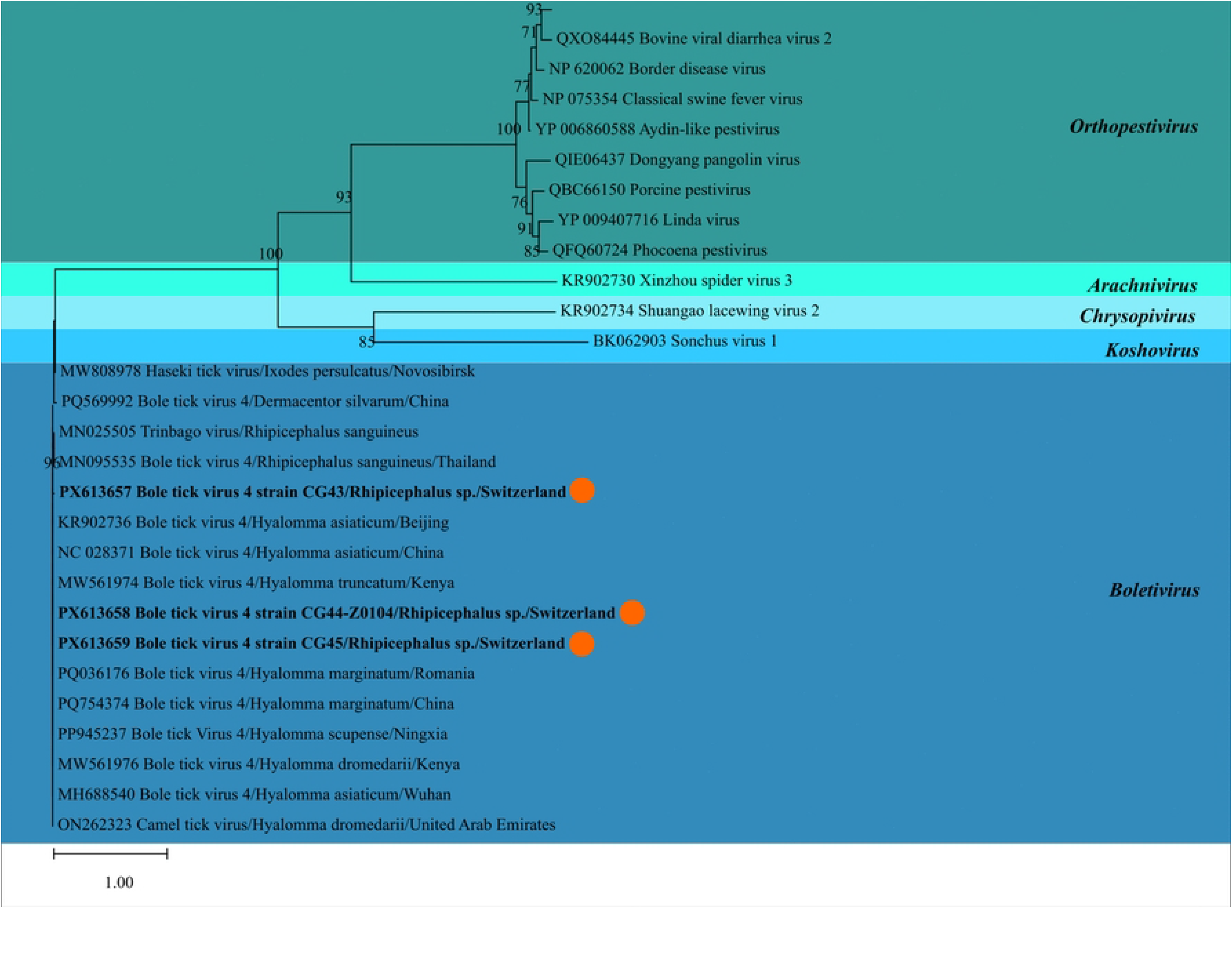
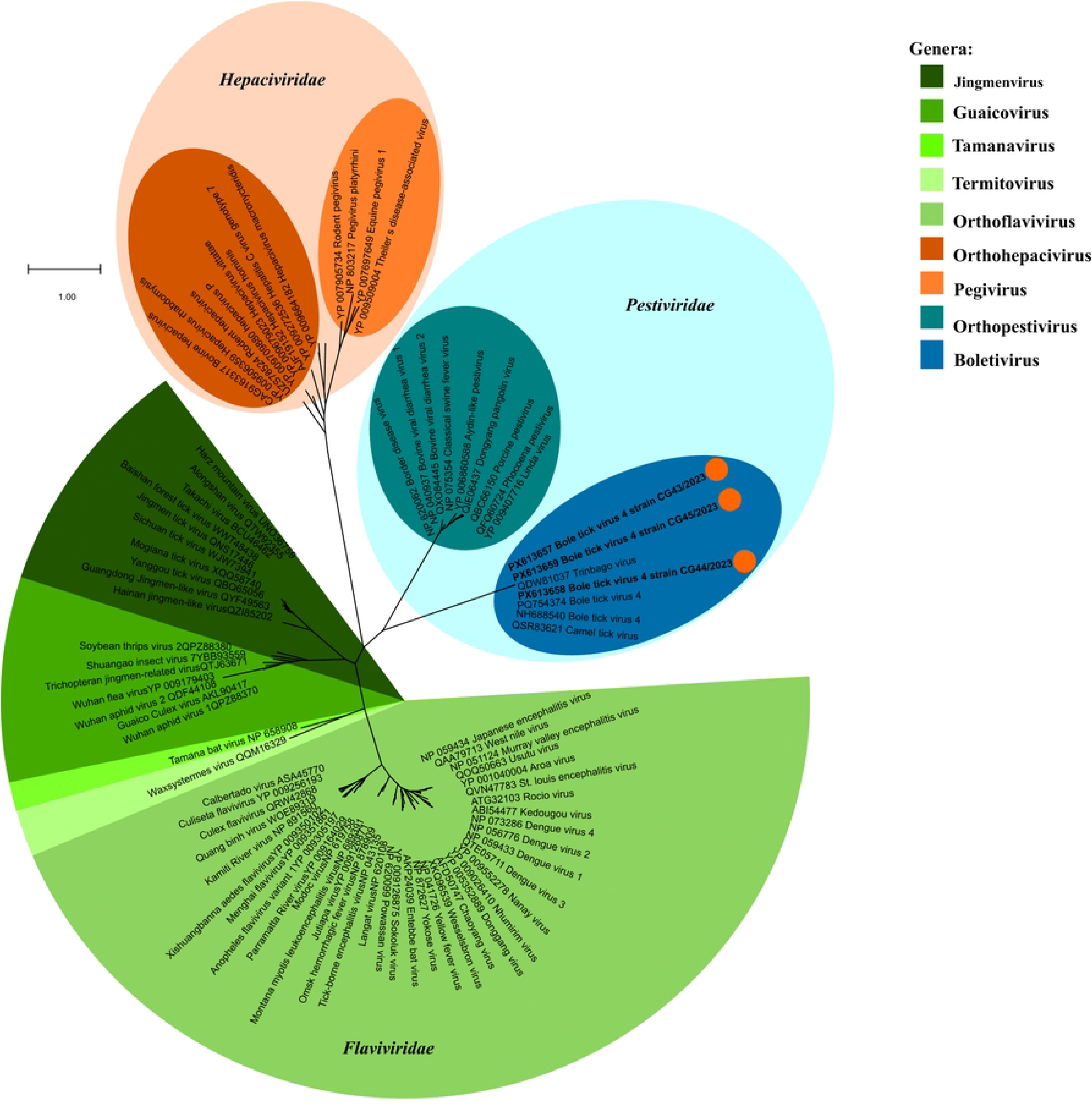
Phylogenetic analysis of Bole tick virus 4 (BoTV4) and related members of the order *Amarillovirale*s. The Maximum Likelihood (ML) tree was constructed using IQ-TREE 2 (v2.3.4) based on an alignment of amino acid (aa) sequences of the RNA-dependent RNA polymerase (RdRp) performed using MAFFT (v7.526), followed by alignment trimming with trimAl (gap threshold: 0.9) to retain highly conserved domains. Numbers at the internal nodes represent Ultrafast Bootstrap (UFBoot2) support values derived from 1,000 replicates. For clarity, only bootstrap values ≥ 70% are displayed. Sequences obtained in this study are indicated in bold and marked with an orange dot (long-distance migratory host bird species). **A.** Analysis of representative members of the order *Amarillovirales,* showing the placement of BoTV4 among the families *Flaviviridae (*including the genus *Orthoflavivirus,* and the newly proposed genera *Jingmenvirus, Guaicovirus, Tamanavirus and Termitovirus), Pestiviridae (*including the genus Orthopestivirus and the newly proposed genus *Boletivirus), and Hepaciviridae (*including the genera *Orthohepacivirus and Pegivirus).* **B**. Detailed analysis of representative members of the family *Pestiviridae*, highlighting the relationships between the genus *Orthopestivirus*, and the newly proposed genera *Arachnivirus, Boletivirus, Chrysopivirus* and *Koshovirus*.

## Discussion

Over a seven-month period, 496 wild birds were sampled, representing three resident and 39 migratory species, ranging from short- to long-distance migrants. Oropharyngeal-cloacal swabs from all birds were initially screened via RT-qPCR for specific viruses, with 476 samples and 22 associated ticks subsequently undergoing metagenomic shotgun NGS. To complement these findings, 164 sera were analyzed by virus-specific ELISA, with all positive or borderline results validated through serum neutralization testing (SNT).

### Host ecology, migration, and seasonal dynamics

The diversity of bird species identified (*n* = 42) and the distribution of sampled individuals *(n* = 496) are closely linked to the ecological function of the sampling site Bolle di Magadino and the sampling strategy. Reflecting seasonal shifts in avian populations, the datasets comprised 268 resident and short-distance migrants and 208 long-distance migrants (Fig 1A and 1B). This distribution is primarily driven by high sampling intensity during the spring migration peak in April and May (S1 Table), when numerous species fly through this area, including long-distance migrants arriving from southeastern Africa via the Po Valley, as well as individuals following South-West routes. During the breeding season (mid-May to end of June), avian movement becomes more localized. Accordingly, the sampled species were primarily territorial residents or breeding migrants and may reflect a relatively stable, so called “local virome” of the Ticino avian population. The post-breeding season is defined by the dispersal of juvenile birds and the arrival of migrants from northeastern routes. The dominance of *Sylvia atricapilla* (*n* = 143) and *Erithacus rubecula* (*n* = 53) in the datasets reflects their status among the most frequent and abundant bird species in Switzerland [92,93]. In a broader context, this representation is of particular significance, according to previous studies *S. atricapilla* has been identified as one of just 16 species contributing to approximately 73% of the billions of birds migrating annually along the Africa-Eurasian flyway [6]. Furthermore, it underscores the importance of the Magadino delta as a major stopover point for rest before continuing migration across the Alps into Central Europe [6,33,52]. From an epidemiological perspective, the site facilitates close intermingling among bird species that are otherwise geographically separated, thereby increasing the potential for interspecies pathogen exchange and dissemination across various migratory flyways [94].

### Molecular and serological results for zoonotic viruses

Zoonotic viruses such as AIV and WNV are of particular concern due to their significant impact on both public health and animal populations [94–96]. In our samples, *Orthomyxoviridae* were detected neither by RT-qPCR nor by NGS. The absence of AIV is most plausibly explained by the host species composition, in which 90.5% of the sampled individuals belonged to the order *Passeriformes*, while key reservoir hosts for AIV, particularly *Anseriformes* (waterfowl) and domestic poultry were not included [45,97,98]. Although some passerines can be experimentally infected with AIV, there is currently insufficient evidence to support their role as natural reservoirs or key transmitters under field conditions [21]. However, sporadic detections in wild passerines, such as the identification of highly pathogenic avian influenza (H5N1) in a dead crow in Kerala (India) in 2026, illustrate that these species can become infected in wildlife [99,100]. Within the *Flaviviridae* family, WNV was a primary target due to its relevance for vector-borne monitoring in Ticino, where the first detection of WNV Lineage 2 in mosquitoes was reported in 2022 [23]. The detection of WNV in mosquitoes represented an early signal of viral circulation and preceded the first confirmed autochthonous human case in Switzerland in 2025 [101]. Birds, particularly *Passeriformes*, are considered highly competent amplifying hosts, frequently developing high viremia levels sufficient to infect mosquitoes. Despite this, no WNV RNA was detected by RT-qPCR or NGS [94,102,103]. However, positive results were obtained in ELISA testing, indicating prior exposure to WNV in the absence of detectable active viremia. This discrepancy between serological and molecular findings is consistent with the observations in Germany (2014–2016), where seropositivity in birds preceded the first autochthonous cases in 2018 despite an initial lack of detectable viral RNA [104]. The absence of viral reads in our unbiased NGS survey, consistent with the negative RT-qPCR results, is likely explained by the biological kinetics of avian WNV infections, where systemic viremia usually lasts only one to eight days. Following the acute phase, viral loads outside the bloodstream decline rapidly, substantially reducing viral detection from swabs, even in recently exposed birds [48,105,106]. The inability to confirm the ELISA positive results using the serum neutralization test (SNT) may be attributed, firstly, to antibody titers falling below the assay detection limit, potentially compounded by limited serum volume available for testing. Secondly, serological cross-reactivity within *Flaviviridae*, specifically among members of the genus Orthoflavivirus, remains a well-recognized limitation. In particular, WNV antibodies exhibit extensive cross-reactivity with other antigenically related viruses, such as Usutu virus (USUV), Japanese encephalitis virus (JEV), and Zika virus (ZIKV), primarily due to highly conserved epitopes within the fusion loop domain of the envelope (E) protein [57,104,107]. In conclusion, while our serological results suggest prior exposure to orthoflaviviruses, the absence of genomic reads via NGS and the negative RT-qPCR results suggest that these findings are more likely explained by survivors infected in other regions who reached Bolle di Magadino after their viremic phase. However, additional spillover events can’t be excluded, including transmission via infected mosquitoes within Ticino or via birds arriving while still viremic [23,105,106,108].

### Virome composition of the birds

The quantitative dominance of small DNA viruses is a recurrent feature of vertebrate viromes, frequently representing the vast majority of identified viral sequences in fecal and swab samples. Consistent with prior metagenomic studies, our detected DNA viruses, dominated by the families *Circoviridae* and *Parvoviridae*, showed lower diversity but accounted for the highest overall viral read count [20,28,37,47,60,109–111], with no significant abundance difference across migratory strategies (Fig 3, Fig 4). This suggests that the core viromic composition remains remarkably stable across migratory strategies, indicating that while migration facilitates pathogen dispersal, the overall swab virome is not primarily shaped by migration. Similarly, while RNA viruses exhibited higher overall family richness (17 families), differential abundance analysis revealed no statistically significant differences in viral abundance across migratory strategies for any RNA viral family (Fig 4). In theory, one might expect long-distance migration to alter virome composition, as long-distance migrants cross multiple ecological zones and are exposed to distinct viral pressures [26,29,44]. In addition, flight-associated metabolic stress during long flights can cause temporary immunosuppression and increased viral shedding, thereby enhancing virus detectability in NGS or in molecular assays, potentially explaining occasional high viral loads in the samples [13,94]. Nevertheless, our results indicate that these individual-level phenomena do not translate into systemic or population-level differences across migratory behaviour. A substantial proportion of the viral families identified in our study is associated with either environmental hosts (e.g., plants and fungi) or invertebrate-infecting viruses (e.g., insects), with most being positive-stranded ssRNA viruses [29,40,112]. The high genetic diversity of these viral groups likely facilitates their widespread distribution across various environmental reservoirs [34,113], aligning with previous research, which consistently detected similar viral composition regardless of the specific avian order analyzed, spanning *Passeriformes, Anseriformes,* and *Charadriiformes* [24,44,46,114]. Overall, the viral composition is strongly influenced by sample type (oropharyngeal-cloacal swabs) and the dietary habits of the host, which likely reflect a mixture of local dietary possibilities and habitat at the sampling site, rather than host species or migratory strategy alone [2]. Consequently, our findings refute the initial hypothesis that the geographic origin of the birds significantly influences the overall virome composition. Instead, the high similarity in environmental and dietary viral signatures across groups indicates that shared exposure at the Bolle di Magadino is the primary determinant of the swab virome. Consequently, many detected viruses may be transient, acquired through ingestion of food, water or through contact with arthropods, rather than indicating active infection in the host [24,60].

### Viral families of interest in the avian virome

#### DNA virome

The metagenomic analysis revealed a quantitative dominance of DNA viruses, particularly from the phylum *Cressdnaviricota* (Fig 5), a pattern frequently observed in vertebrate metaviromics [24,60]. This phylum comprises small circular ssDNA viruses, including the *Circoviridae* and *Genomoviridae*, belonging to the group known as CRESS-DNA viruses (circular Rep-encoding single-stranded DNA viruses), which are characterized by genomes ranging from 1 to 25 kb that encode a Rep protein to initiate replication via a rolling-circle mechanism [115–117]. While birds are recognized as important hosts for these viruses, the full extent of their circular ssDNA diversity remains poorly characterized. Notably, our three CRESS-DNA contigs did not cluster with known avian pathogens, a finding consistent with large-scale research from China, which has shown that *Passeriformes* serve as an important natural reservoir for highly diverse CRESS-DNA lineages, often forming divergent clades related to environmental or vector-associated viruses [24]. Furthermore, although CRESS viruses have been detected in human clinical samples, their pathogenic relevance remains unconfirmed [118]. This DNA dominance is further reflected in the family *Parvoviridae*, which was among our most abundant and diverse groups. The viruses of this family are frequently present in the avian intestinal tract, often found in healthy and diseased birds [25,119–121]. Interestingly, the three *Parvoviridae* contigs identified in this study were all classified within the subfamily *Densovirinae* (Fig 6). Given that Densoviruses primarily infect arthropods, their presence likely reflects dietary acquisition of insects rather than an active vertebrate infection. The host composition of our study is directly reflected in the low abundance or absence of certain viral families that are otherwise ubiquitous in avian metaviromics. While *Adenoviridae* and *Caliciviridae* are considered global residents in wild birds, they were present at low abundance or absent in our samples [26,112,114,122–125]. This distinct viral profile may be attributed to the lack of aquatic birds (*Anseriformes* and *Charadriiformes*) that typically serve as major reservoirs for many of these absent families [2]. Finally, members of the *Microviridae* were identified despite nuclease treatment, representing a highly diverse group of circular ssDNA bacteriophages previously described in the gut virome of mammals and birds, particularly within the order Passeriformes [126,127].

#### RNA virome

Avian gastrointestinal viromes are generally characterized by a core group of viral RNA families, typically dominated by the *Picornaviridae,* which often account for approximately 30% of viral reads in avian fecal and gut samples [20,119,128]. Our data, however, showed a low prevalence, suggesting that the local virome at the Bolle di Magadino is primarily shaped by site-specific biodiversity and diet rather than a stable core of vertebrate viruses. This ecological signature is further evidenced by the presence of invertebrate-infecting virus families (Fig 7), such as *Dicistroviridae*, *Iflaviridae*, and *Polycipiviridae*, which likely represent transient “passengers” acquired through arthropod ingestion rather than active vertebrate infection [26,129–131]. While our findings for *Dicistroviridae* align with prevalences reported in migratory populations across China, the contrastingly lower levels of Iflaviridae likely reflect the specific local composition and prey availability at the Bolle di Magadino site [26,29,114]. Beyond their role as dietary markers, these findings highlight the potential role of migratory birds as incidental mechanical vectors of agriculturally relevant pathogens across different ecological zones [13,132]. The birds’ close interaction with the local ecosystem is further exemplified by viruses with broader environmental associations. This includes the detection of *Orthototivirida*e, which encompassed contigs linked to both insect and plant hosts [133]. Together with the presence of plant-infecting families such as *Alphaflexiviridae* and *Bromoviridae*, these findings reflect an exposure profile resulting from direct consumption of infected material or insects and contact with the surrounding vegetation at the Bolle di Magadino site [114,134,135].

#### Detection of Bole tick virus 4 (BoTV4) in ticks

Within the family *Pestiviridae*, we identified Bole tick virus 4 (BoTV4) in ticks of the species *Rhipicephalus turanicus*, marking its first detection in Switzerland. Initially described in 2015 through metagenomic sequencing in China, BoTV4 has subsequently been reported in multiple tick genera, predominantly *Hyalomma* and *Rhipicephalus*, across Asia, Europe, Africa and the Caribbean [91,136–142]. Historically, the family *Pestiviridae* was accommodated as a lower-rank taxon (the genus Pestivirus) within the family *Flaviviridae*, alongside the genera *Orthoflavivirus, Hepacivirus*, and *Pegivirus*. However, under the major 2025 taxonomic reorganization of the order *Amarillovirales*, these classical lineages have been elevated to distinct family-level ranks: *Flaviviridae* (Clade I), *Pestiviridae* (Clade II), and *Hepaciviridae* (which also includes the genus *Pegivirus*, Clade III). The expanded family *Pestiviridae* absorbs the former genus *Pestivirus* (now renamed *Orthopestivirus*) but also encompasses highly divergent arthropod-associated lineages formerly grouped under the "large-genome flaviruses" (LGFs), of which BoTV4 is a prominent member officially assigned to the newly established genus Boletivirus [89]. BoTV4 distinguishes itself from classical members of the *Flaviviridae* by possessing an enlarged, non-segmented, positive-sense ssRNA genome of approximately 16.3 kb, encoding a single large polyprotein [138,140,141]. While BoTV4 shares conserved non-structural protein (NS3 and NS5) with the Jingmenvirus (JMV) group (e.g., Alongshan virus and Jingmen tick virus) belonging to Clade I, it lacks their characteristic genomic segmentation [90,143]. Instead, BoTV4 represents a unique evolutionary lineage. Our sequences fall firmly within Clade II, clustering entirely separate from both the classical vector-borne and insect-borne *Flaviviridae* und mammalian *Hepaciviridae* (Fig 8A). On family-level, the newly characterized Swiss strains form a highly supported, separate monophyletic clade officially designated as the genus Boletivirus (Fig 8B). While the replication mechanisms of BoTV4 and its zoonotic potential remain incompletely understood, sequences of this virus and its close relatives e.g., Haseki tick virus have been detected in human serum samples from febrile patients [138]. Furthermore, metagenomic analysis of the skin infectome at the tick bite site has identified sequences related to BoTV4, often associated with localized clinical signs such as papule at the site of attachment [144]. The detection of BoTV4 in ticks collected from *A. arundinaceus* at the Bolle di Magadino highlights the potential role of migratory birds, particularly long-distance migrants, in the dispersal of ectoparasites and their associated pathogens. These ticks may have acquired in Mediterranean or sub-Saharan regions prior to flight into Switzerland. In our dataset 22 ticks were collected from 496 birds, representing only a small fraction relative to the scale of avian migration across the Palearctic-Afrotropical flyways, where more than two billion birds migrate annually between Europe and Africa [6]. Migratory birds can therefore act as mechanical dispersal hosts, transporting attached ticks across continental borders and ecological barriers within days [145]. Consequently, the detection of BoTV4 highlights the need for continued surveillance of avian-associated ectoparasites to better assess the introduction and circulation of emerging or novel tick-borne viruses in Switzerland.

## Conclusion

This study represents the first active monitoring of wild bird virome in Switzerland, providing a comprehensive investigation of oropharyngeal-cloacal samples from 496 birds (42 species) and 22 associated ticks. We identified 29 distinct viral families in the birds, expanding the understanding of avian viral diversity. While no emerging or exotic viruses were identified within the avian viromes, our results support the hypothesis that migratory birds may increase the risk of introducing novel pathogens into Switzerland by transporting infected ectoparasites, as demonstrated by the first detection of Bole tick virus 4 (BoTV4, family *Pestiviridae*) in ticks collected from long-distance migrants. Overall, the composition of the avian virome primarily reflects the local ecological interaction, habitat, and dietary behaviour. The majority of detected viruses appear to be transient, reflecting environmental exposure rather than active infection, and no evidence supporting zoonotic potential in the avian virome was identified here [26,114]. However, the positive WNV ELISA results indicate that some birds had previous exposure to *Orthoflaviviruses*, representing survivors of an earlier infection phase. While these seropositive birds may have reached the Bolle di Magadino after their acute viremia, it is noteworthy that the viremic phase in passerines typically lasts between five and seven days [108]. This duration is sufficient for individuals to reach subsequent stopover sites while still infectious. Once introduced at a new location closer to Switzerland, the virus can establish a local transmission cycle, thereby contributing to the step-by-step dispersal of virus across ecological zones [12]. However, it should be noted that while swab-based metagenomics is highly effective for revealing diversity, it provides limited sensitivity for detecting systemic infections at low viral loads. In this context, although limited and more challenging, blood-based virome profiling may be necessary to capture more comprehensively active infections [146]. Taken together, our results highlight the importance of continued monitoring of not only migratory birds but also bird-associated ectoparasites, particularly ticks, to improve early detection and monitoring of emerging tick-borne threats.

## Acknowledgments

We thank the Next Generation Sequencing Platform of the University of Bern for performing the high-throughput sequencing experiments. We sincerely thank Marie-Anne Berazategui and Kin Morandi for their invaluable contribution to the collection of bird samples. We are also grateful to collaborators of the Fondazione Bolle di Magadino and Ficedula, in particular Nicola Patocchi, Roberto Lardelli, and Chiara Scandolara, for their dedicated support in bird sampling activities. Furthermore, we thank the Institute of Parasitology at the University of Bern, specifically Walter Basso and Maryna Galat, for their expert contribution to the morphological characterization of the ticks.

## Supporting Information

S1 Table. Master metadata, sample details, metagenomic sequencing statistics, and database accession numbers for wild birds and associated ticks in Ticino, Switzerland (2023). (XLSX)

S2 Table. Individual serological screening results for Orthoflavivirus antibodies and WNV serum neutralization in sampled wild birds (Ticino, 2023). 28 individual birds with positive/doubtful ELISA results were tested by WNV SNT. (DOCX)

